# Single cell transcriptomic landscapes of human liver organoids stratify models of non-alcoholic fatty liver disease

**DOI:** 10.1101/2022.07.19.500693

**Authors:** Anja Hess, Stefan D. Gentile, Amel Ben Saad, Raza-Ur Rahman, Tim Habboub, Alan C. Mullen

## Abstract

Non-alcoholic fatty liver disease (NAFLD) is a rapidly growing cause of morbidity with few treatment options available. Thus, accurate *in vitro* systems to test new therapies are indispensable. Recently, human liver organoid (HLO) NAFLD models have emerged. However, a systematic evaluation of their translational potential is currently missing. Here, we develop a structured approach to evaluate NAFLD-HLO models, testing oleic acid (OA) and palmitic acid (PA) in comparison to TGF-β1 for disease induction. Through analysis of ∼100K single-cell transcriptomes of the HLO injury landscape, we find all three models induce inflammatory signatures. However, only TGF-β1 promotes collagen production, fibrosis, and hepatic stellate cell (HSC) expansion. In striking contrast, OA ameliorates fibrotic signatures and reduces the HSC population. Integrating data from each model with that of NAFLD patients across disease progression further demonstrates PA and TGF-β1 more robustly model inflammation and fibrosis. Our findings highlight the importance to stratify NAFLD-HLO models by clinical disease progression, provide a single-cell reference to benchmark future organoid injury models, and allow us to study evolving steatohepatitis, fibrosis, and HSC susceptibility to injury in a dynamic, multi-lineage human *in vitro* system.

## Introduction

Chronic liver injury promotes sustained inflammation, leading to liver fibrosis, which can progress to cirrhosis^1^, a major cause of morbidity and mortality worldwide^2^. Non-alcoholic fatty liver disease (NAFLD) is among the most common causes of chronic liver disease^2^ and the most rapidly increasing indication for liver transplantation in the US^3^. NAFLD is tightly linked to obesity and metabolic syndrome^4^. While the majority of NAFLD cases that lead to end-stage liver disease progress from simple steatosis (nonalcoholic fatty liver, NAFL) to non-alcoholic steatohepatitis (NASH, inflammation due to steatosis) and then fibrosis^5,6^, it is important to acknowledge that NAFL is a prevalent finding in otherwise healthy individuals. Indeed, only approximately 30% of individuals with NAFL develop NASH^7^. While the progression from NASH to fibrosis is well documented, there are currently few treatment options to disrupt this process other than weight loss^8^. More broadly, there are no approved treatments available that target common inflammatory or fibrotic pathways arising from chronic liver injury, which could prevent the progression of NASH or other chronic liver diseases^8^. Thus, the development and evaluation of *in vitro* human cell models of liver inflammation and fibrosis is critical for creating and testing new approaches to prevent liver failure.

A key prerequisite for NAFLD-associated chronic liver injury is the continued exposure to excess fatty acids^9–11^. The free fatty acids (FFAs) oleic acid (OA) and palmitic acid (PA) accumulate in NAFLD and thus are widely used to model NAFLD *in vitro*^12,13^. OA, a monounsaturated fatty acid (MUFA), induces hepatocyte steatosis but has also been reported to exert protective effects against NAFLD^14,15^. PA, a saturated fatty acid (SFA), promotes hepatocyte apoptosis^16,17^.

FFA-induced hepatocyte damage activates pro-inflammatory signaling pathways in resident liver cells^4^. This leads to the secretion of cytokines such as tumor necrosis factor (TNF)^4,5^, activating resident liver macrophages (Kupffer cells)^18^. Kupffer cells and recruited immune cells^19^ produce transforming growth factor beta (TGF-β)^20^, which activates hepatic stellate cells (HSCs), and promotes their differentiation towards myofibroblasts^21^. These HSC myofibroblasts produce extracellular matrix (ECM, predominantly collagen type I and III) that accumulates to form the fibrotic scar^21^. Hallmarks of fibrosis include the up-regulation of transcripts *COL1A1, COL3A1, TIMP1*, *TNFS* members and the receptor-ligand-pair *PDGFB/PDGFR*^21,22^. Additionally, chronic liver injury is also associated with the expansion of cells with ductular characteristics, which can originate from cholangiocytes, hepatic progenitors, and hepatocytes to replace injured hepatocytes^23–26^.

Recently, the generation of multi-lineage hepatic organoids from human pluripotent stem cells (hPSCs) has emerged^27–31^. These systems contain cells of at least endodermal- and mesodermal identity (e.g., hepatocyte-, cholangiocyte-, and hepatic stellate cell-like cells) and are also referred to as multi-tissue organoids^32^. Some multi-lineage organoids, herein referred to as human liver organoids (HLOs)^27^, have been shown to recapitulate aspects of liver inflammation and fibrosis^27,28^, and thus are promising *in vitro* models for NAFLD. However, to validate their translational potential, a comparative evaluation of such systems is urgently needed. Single-cell RNA sequencing (scRNA-seq) has emerged as a key tool to identify disease signatures in human organoids at high resolution^33^. To our knowledge there are currently no studies comparing the hepatic injury type and NAFLD severity induced by different agents in HLO models at single-cell resolution.

Here, we develop a structured approach to evaluate NAFLD models in an HLO system and provide a reference of ∼100,000 single-cell transcriptomes reflecting the HLO injury landscape. We examine OA, PA, and TGF-β1 for their potential to induce collagen production and select the optimal 3D culture environment for this purpose. Next, we apply functional and 10X scRNA-seq transcriptomic analysis to evaluate the induction of inflammatory and fibrotic signatures. Finally, we apply clinical NAFLD gene signatures to score the severity of the generated HLO injury. This approach allows us to stratify *in vitro* NAFLD models by their alignment to disease progression in patients and identifies PA as the more robust FFA model of NASH.

## Results

### Human liver organoids recapitulate the transcriptional landscape of major cell types in the adult human liver

We first differentiated hPSCs into HLOs as described^27,34^ (**Fig. 1a**). We confirmed loss of the canonical pluripotency genes *SOX2*, *NANOG,* and *POU5F1* by day 16 of differentiation and induction of the human liver genes *ASGR1* and *HNF4A* (hepatocyte markers^35^), *KRT19/CK19* and *SOX9* (cholangiocyte markers^36^) as well as *VIM* and *DES* (HSC markers^37^) in day 21 HLOs (**Fig. 1b**). To evaluate HLO identity at the protein and histologic level, we performed hematoxylin and eosin (H&E) staining and immunohistochemistry (IHC) and found that HLOs at day 21 display an organized, sphero-luminal structure, and multiple cells express the nuclear hepatocyte marker CEBPα (**Fig. 1c**, Supplementary Fig. 1). These results are in agreement with previous observations on the emergence of distinct liver cell types from at least two different lineages in HLOs after day 20^27^.

**Fig. 1.**
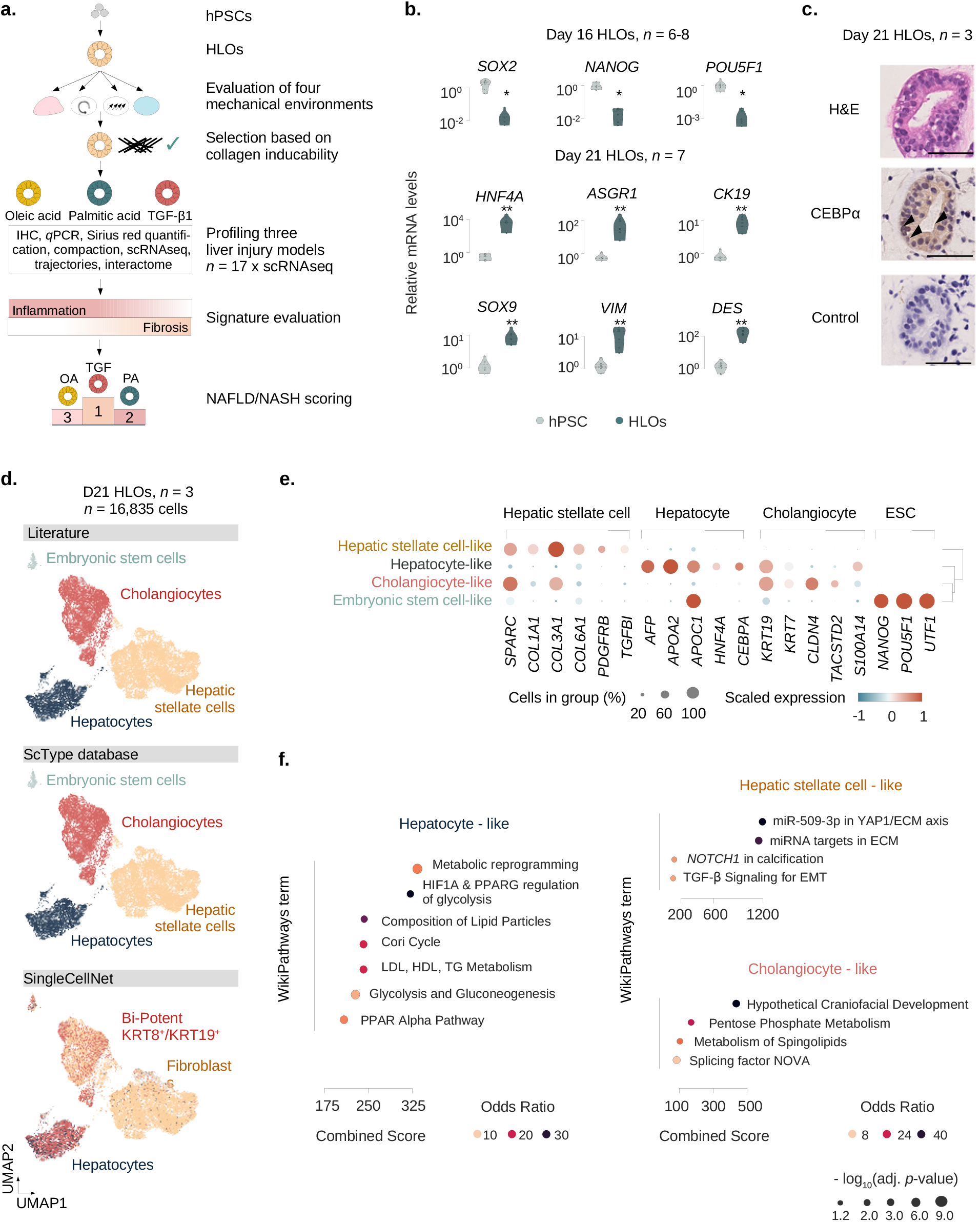
Human liver organoids recapitulate the transcriptional landscape of major cell types in the adult human liver. **a.** Overview of the experimental design. Human pluripotent stem cells (hPSCs) are differentiated into human liver organoids (HLOs) and cultured in four different mechancial environments. The environment suitable for collagen induction is selected, and three injury scenarios (OA, PA, TGF-β1) are evaluated for their potential to model inflammation and fibrosis by functional and transcriptomic readouts, with a total of 17 HLO samples being subjected to 10X single-cell RNA sequencing (scRNA-seq). Transcriptomic signatures for inflammation and fibrosis are derived and evaluated. Finally, a NAFLD/NASH severity score is applied, allowing for the hierarchical ordering of HLO injury models by their potential to model NAFLD disease progression. **b.** Expression of pluripotency genes is reduced with HLO differentiation, while liver markers are induced. Relative gene expression was measured by quantitative reverse transcriptase (qRT) PCR and normalized by housekeeping gene *ACTB* and displayed relative to hPSC controls (2^-ddCt^). Comparison between hPSCs and HLOs from the same experiment were performed at day 16 or 21 as indicated. *N* = 6-8 individual experiments as indicated by circles. Mann-Whitney-U-statistics (two-tailed) hPSCs vs. HLOs: *p*-values^day 16 HLOs^ = 0.0024, Bonferroni-adjusted level of significance: 0.0125 (*); *p*-values^day 21 HLOs^ = 0.0022, Bonferroni-adjusted level of significance: 0.0071 (**). **c.** HLOs on day 21 show an organized, luminal structure and express human liver proteins. Representative immunohistochemistry images of HLOs stained in hematoxylin & eosin (H&E), CEBPα and negative control with secondary antibody on day 21. Arrowheads indicate sites of nuclear protein staining. Scale bars, 50 µm. *N* = 3 individual experiments. **d.** scRNA-seq analysis of day 21 HLOs identifies the major cell types of the human liver across three different annotation strategies. UMAP plots showing cells from 10X scRNA-seq in day 21 HLOs (*n* = 3 replicates). Cell type annotations generated by three different methods are displayed: Literature-based annotation (top), ScType^40^ database (center), and SingleCellNet^42^ annotation comparing HLOs to human liver single-cell data^41^ (bottom). **e.** Dotplot showing the expression of canonical cell-type-specific marker genes expected in the human liver across clusters. Cell types defined by single-cell data are displayed on the y-axis, and marker genes (bottom) are sorted by cell type (top). The fraction of cells expressing a gene is indicated by the size of the circle, and the scaled mean expression of a gene is indicated by color. Hierarchical clustering is represented by the dendrogram on the right. **f.** Top enriched WikiPathways^82^ terms for the three major cell types in day 21 HLOs. Terms are sorted by combined score and have been shortened for readability. Dot sizes correspond to the negative decadic logarithm of the adjusted *p*-value, dot color represents the Odds ratio for each term.

To further investigate the cellular composition of HLOs, we performed 10X scRNA-seq on day 21 and analyzed a total of 16,835 cells after quality control (QC) (**Fig. 1d**, Supplementary Fig. 2, Methods). Annotation of HLO populations can create difficulties since differentiating systems may contain transient cell states and retain premature features^38^. To address these challenges, we initially evaluated three different annotation strategies^39^. We first annotated cell clusters based on marker genes from the literature for fetal and adult liver cell types (Supplementary Table 1, Supplementary Fig. 2), rendering four main clusters of hepatocyte-like, HSC-like, cholangiocyte-like cells, and a small fraction of embryonic stem cell (ESC)-like cells (**Fig. 1d**, top). In a second approach, we took advantage of a recently published marker gene database validated on human adult liver scRNAseq data (ScType)^40^. This analysis showed similar results to the literature approach (**Fig. 1d**, center, Supplementary Fig. 2). We then evaluated a third approach, applying a random forest-based classifier to compare HLOs to scRNA-seq data from fetal and adult liver tissue^41^. We chose SingleCellNet (SCN)^42^ as it has been reported to yield high accuracy for benchmarking data of human pancreatic cell populations^43^, which developmentally are closely related to the liver^44^. The SCN hepatocyte and HSC-annotations are consistent with the previous methods. Cholangiocytes are mainly assigned to the *KRT8*+/*KRT19*+ bi-potent population, labeled according to the terminology in the reference study^41^ (**Fig. 1d**, bottom, Supplementary Fig. 2e-f). This population is specific to normal, non-tumor adult liver tissue and has been related to cholangiocytes^41^ based on another scRNAseq study of adult human liver^22^.

Overall, the attributions of hepatocyte, HSC, and cholangiocyte identities are overlapping across annotation strategies, and two out of three strategies also indicated a small embryonic-stem cell population as a fourth cell type in day 21 HLOs. Based on this comparative analysis we decided to utilize the ScType^40^ database annotation method for all subsequent analyses since it allows for the annotation of potentially emerging cell types beyond the repertoire of a literature list or a single reference study. To account for the *in vitro* generation of the cells, we refer to them as cell type*-like* in this study. To further ensure cellular identities, we confirmed the expression of canonical marker genes for each cell type and found they overlap with the consensus annotations (**Fig. 1e**). These results support the annotations from the previous strategies, however, *KRT19* was still positive in the hepatocyte-like population, indicating they retained premature features on day 21 in HLOs. We then performed pathway enrichment analysis, revealing upregulated signatures for liver-characterizing metabolic processes and lipid metabolism in hepatocyte-like cells. HSC-like cells were enriched for pathways related to extracellular matrix and focal adhesion. Cholangiocyte-like cells showed enrichment for pentose phosphate metabolism and sphingolipids (**Fig. 1f**). Together, these results indicate the presence of hepatocyte-, HSC-, and cholangiocyte-like cells in day 21 HLOs across cell type annotation strategies.

### Inducible liver injury phenotypes in HLOs upon TGF-β1 and fatty acid treatment

We next evaluated conditions to induce a fibrotic phenotype in the HLO model. TGF-β1 drives liver fibrogenesis *in vivo*^45,46^, and we treated HLOs with TGF-β1 (10 ng/ml and 25 ng/ml) in four different 3D culture systems and evaluated collagen expression. The final differentiation steps for HLOs are performed in Matrigel, and HLOs were either left in Matrigel or removed from Matrigel and cultured on i) ultra-low attachment plates (commercially available cell culture plates coated with a hydrogel layer to avoid attachment) as previously described^27^, ii) 1% agarose coated plates, and iii) an orbital shaker (**Fig. 2a**).

**Fig. 2.**
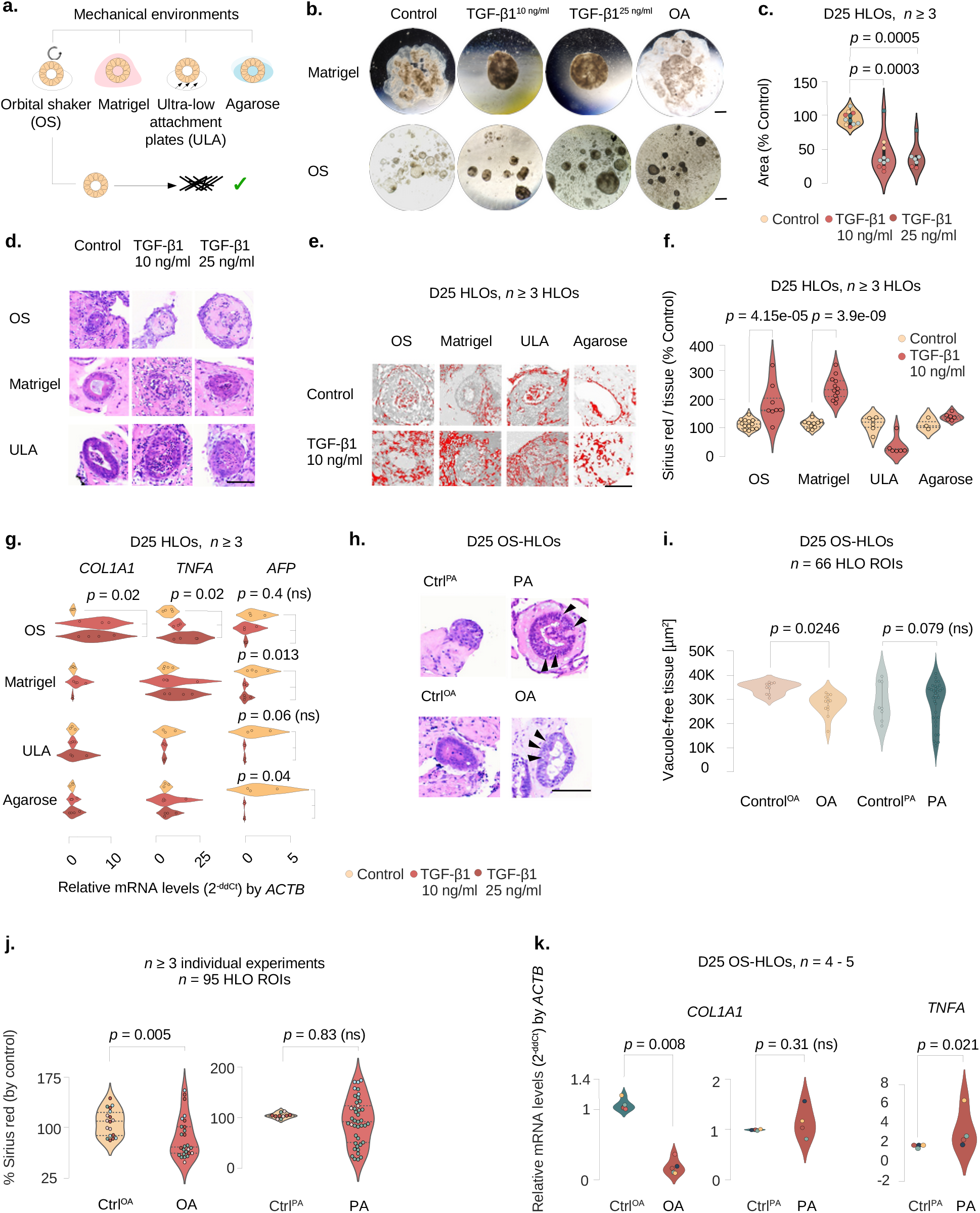
Inducible liver injury phenotypes in HLOs upon FFA and TGF-β1 treatment. **a.** Overview of the four mechanical environments evaluated for their potential to promote collagen induction with injury models. **b.** HLOs in the Matrigel- and OS-environments display morphologic changes when treated with TGF-β1 and OA. Brightfield images of whole Matrigel domes with embedded HLOs (upper panel, scale bar 3 mm) and HLOs isolated from Matrigel in the orbital shaker environment (lower panel, scale bar 0.1 mm) after four days of treatment with TGF-β1 (10, 25 ng/ml), OA (400 µM), and controls. Contraction of the Matrigel dome with TGF-β1 treatment and darkening of color (among all treatments) is observed. **c.** Compaction assay for Matrigel-environment cultured HLOs indicates a significant surface area reduction for HLOs treated with TGF-β1. Violin plots displaying the Matrigel drop area normalized by the control. Kruskal-Wallis test (two-tailed) followed by a post hoc Conover’s test with Bonferroni correction. Dot colors represent experiments, circles represent individual biological replicates. *N* = 3 - 4 experiments. **d.** H&E stainings of OS-HLOs (upper row), Matrigel-HLOs (middle row), and ULA-HLOs (lower row) after four days of treatment with TGF-β1 (10, 25 ng/ml) display wall thickening in Matrigel- and ULA-HLOs while hyaline-like intra-organoidal mass accumulation is present only in OS-HLOs. Scale bar 100 µm. *N* = 1 - 3 experiments. **e.** Thresholded images of Sirius red stained HLOs cultured in four different environments stimulated with TGF-β1 (10 ng/ml), showing collagen deposition in Matrigel- and OS-HLOs. Scale bars 100 µm. *N* ≥ 3 HLOs. **f.** Sirius red staining quantification indicates significant collagen fiber deposition only in HLOs exposed to Matrigel- and OS-environments during stimulation with TGF-β1 (10 ng/ml). Violin plots showing the percentage of Sirius red positive tissue normalized by the mean of the control. Kruskal-Wallis (two-tailed) test followed by a post hoc Conover’s test with Bonferroni correction. *N* ≥ 3 HLOs. **g.** qPCR for *COL1A1, TNFA,* and *AFP* indicates a significant induction of *COL1A1* and *TNFA* transcripts only in HLOs stimulated in the OS-environment. Violin plots showing relative mRNA levels normalized to *ACTB* for each mechanical environment (2⁻^ddCt^). Kruskal-Wallis (two-tailed) test followed by a post hoc Conover’s test with Bonferroni correction comparing three treatment groups per culture method. *P*-values as indicated in the figure, ns, not significant. **h.** HLOs cultured in the OS-environment acquire intracellular vacuoles consistent with steatosis after the exposure to FFAs. H&E stainings of day 25 HLOs treated with PA or OA for four days. Arrowheads indicate vacuoles. Scale bar, 100 µm. **i.** Tissue quantification for 200 x 200 µm regions of interest (ROIs) around single HLOs treated with OA, PA, and their respective controls. Kruskal-Wallis test (two-tailed) followed by a post hoc Conover’s test with Bonferroni correction. *P*-values as indicated in the figure, ns, not significant. *N* = 66 HLOs, *n* = 3 experiments. **j.** Sirius red quantification in HLOs treated with OA, PA, and their controls. Shown is the percentage of Sirius red positive tissue by total tissue (normalized by the mean of the control HLOs in each experiment). Quantification for 200 x 200 µm regions of interest (ROIs) around single HLOs. Kruskal-Wallis test (two-tailed) followed by a post hoc Conover’s test with Bonferroni correction, ns, not significant. *N* = 95 HLOs, *n* ≥ 3 experiments. **k.** *COL1A1* expression is reduced in OS-HLOs treated with OA for four days, *TNFA* expression is induced in OS-HLOs treated with PA for the same duration. Shown are qPCR results for relative mRNA levels of *COL1A1* and *TNFA* normalized to *ACTB*. Mann-Whitney-U test (two-tailed). *P*-values as indicated in the figure, ns, not significant. *N* = 4-5 experiments (represented by individual dot colors).

We observed morphological changes in HLOs cultured in Matrigel and on the orbital shaker after four days of treatment including HLO tissue consolidation, along with surface roughening in TGF-β1-treated HLOs (**Fig. 2b**, Supplementary Fig. 3a). For comparison, we also evaluated HLOs treated with OA, which did not demonstrate the same compaction in Matrigel as observed with TGF-β1 but was associated with a darker appearance on light microscopy when cultured in the orbital shaker, as previously described^27^. These results suggest HLOs are reacting in an injury-specific manner to the applied treatments.

We further quantified the contractile effect of TGF-β1 by culturing HLOs in Matrigel drops and measuring the Matrigel drop area after TGF-β1 application. This analysis demonstrates a significant reduction in droplet size with TGF-β1 treatment, consistent with increased contractile activity (**Fig. 2c**). To further investigate alterations in HLOs we stained for H&E and observed the intraluminal accumulation of hyaline, monomorph structures upon TGF-β1 treatment (**Fig. 2d**), providing further evidence for an injury response to TGF-β1 treatment.

We next performed Sirius red staining (**Fig. 2e**), a standard method for quantification of type I and III collagen deposition in liver fibrosis^47^. We optimized an existing pipeline (Methods) for the computational quantification of Sirius red staining in liver for sections with low tissue amounts and options to select individual HLO areas of interest for the calculation of Sirius red percentage per tissue and per area separately (Methods, github.com/anjahess/sr_organoids). We utilized our pipeline to analyze HLOs cultured via the four previously described methods and treated with TGF-β1 (10 ng/ml) for four days. We found a significant increase of collagen deposition only in Matrigel and orbital shaker-cultured HLOs (**Fig. 2f**), suggesting the latter culture methods lead to accumulation of type I and III collagen as observed in liver fibrosis.

We next evaluated gene expression response across the four culture methods in HLOs when treated with TGF-β1. Canonical transcriptional changes in steatohepatitis include the activation of *TNFA*^1^, while induction of the alpha-1 subunit of type 1 collagen (*COL1A1*) is characteristic of fibrosis^22^. We analyzed mRNA levels for the respective genes. We found that only HLOs cultured on an orbital shaker with TGF-β1 showed a significant increase in *TNFA* and *COL1A1* expression (**Fig. 2g**), while *AFP* was reduced in HLOs cultured on Matrigel and 1% agarose, potentially reflecting different levels of TGF-β1-responsive progenitor-like cells across culture conditions^48,49^. These results show that culturing in the orbital shaker provides the conditions under which HLOs demonstrate the most robust inflammatory and fibrotic response to TGF-β1.

We therefore focused on the orbital shaker method to evaluate the response to fatty acids. Treatment with OA and PA resulted in the formation of lipid droplets, observed as stain-free vacuoles on H&E (**Fig. 2h**, arrowheads), and we assessed lipid accumulation by quantifying the proportion of stain free tissue in regions of interest (ROIs, 200 x 200 µm) capturing individual organoids. Only OA-treated HLOs showed significant tissue rarefaction due to steatosis (**Fig. 2i**, Supplementary Fig. 3b). We then sought to understand if OA and PA could induce collagen production. Surprisingly, OA treatment led to a reduction in Sirius red staining and the quantified percentage of Sirius red positive tissue, as a marker of collagen deposition, while PA did not change Sirius red staining (**Fig. 2j**, Supplementary Fig. 3c). In line with these results, treatment with OA significantly reduced *COL1A1* mRNA levels at the transcriptional level that did not change with PA (**Fig. 2k**). However, PA treatment led to a significant increase in *TNFA* mRNA levels, whereas OA was associated with a significant reduction in *TNFA* at 400 µM concentration (**Fig. 2k**, Supplementary Fig. 3d). These results show that OA, but not PA, induced significant steatosis, while PA but not OA induced *TNFA* expression consistent with an inflammatory response. Neither OA nor PA induced fibrotic injury, with OA treatment actually reducing levels of collagen at both the transcriptional and protein level.

### Hepatocyte maturation and cell type distribution are regulated by mechanical culture conditions

The induction of *COL1A1* mRNA in response to TGF-β1 was only observed in HLOs cultured on an orbital shaker (OS-HLOs, see for Figs. 2d-g). To gain insights into the differences between OS-HLOs and HLOs conventionally cultured on an ultra low attachment plate with 10% Matrigel^34^ (ULA-HLOs), we analyzed a total of 11 HLO samples on ULA plates (*n* = 3 ULA day 21, Fig. 1d) and control HLOs on the OS (*n* = 8 from day 25) (**Fig. 3a**). After joint QC and normalization, our dataset contained a total of 16,835 single cells from ULA-HLOs and 49,011 single cells from OS-HLOs. We next ran our standard pipeline, which detected four cell clusters in each HLO culture context and projected the clusters on a force-directed layout for optimal global structure preservation^50^. ULA-HLO cells displayed the same cell types shown in Fig. 1d, and the OS-HLOs were annotated as hepatocyte-, cholangiocyte-, HSC-, and fibroblast-like cells (**Fig. 3b**). No embryonic stem cell-like cells were present in OS-HLOs. We further observed that ULA-HLOs contained a greater proportion of cholangiocyte-like cells (**Fig. 3c**). Additionally, the ratio of hepatocyte-like to HSC-like cells was higher in OS-HLOs when compared to ULA-HLOs. We also confirmed the broader distribution of HSC marker genes in ULA-HLOs (**Fig. 3d**). Together these results indicate that OS-HLOs show a distribution of cell types that more closely resembles the adult liver compared to ULA-HLOs. The relative reduction of HSCs in OS conditions may also create more room for expansion of HSCs and *COL1A1* induction with TGF-β1 treatment.

**Fig. 3.**
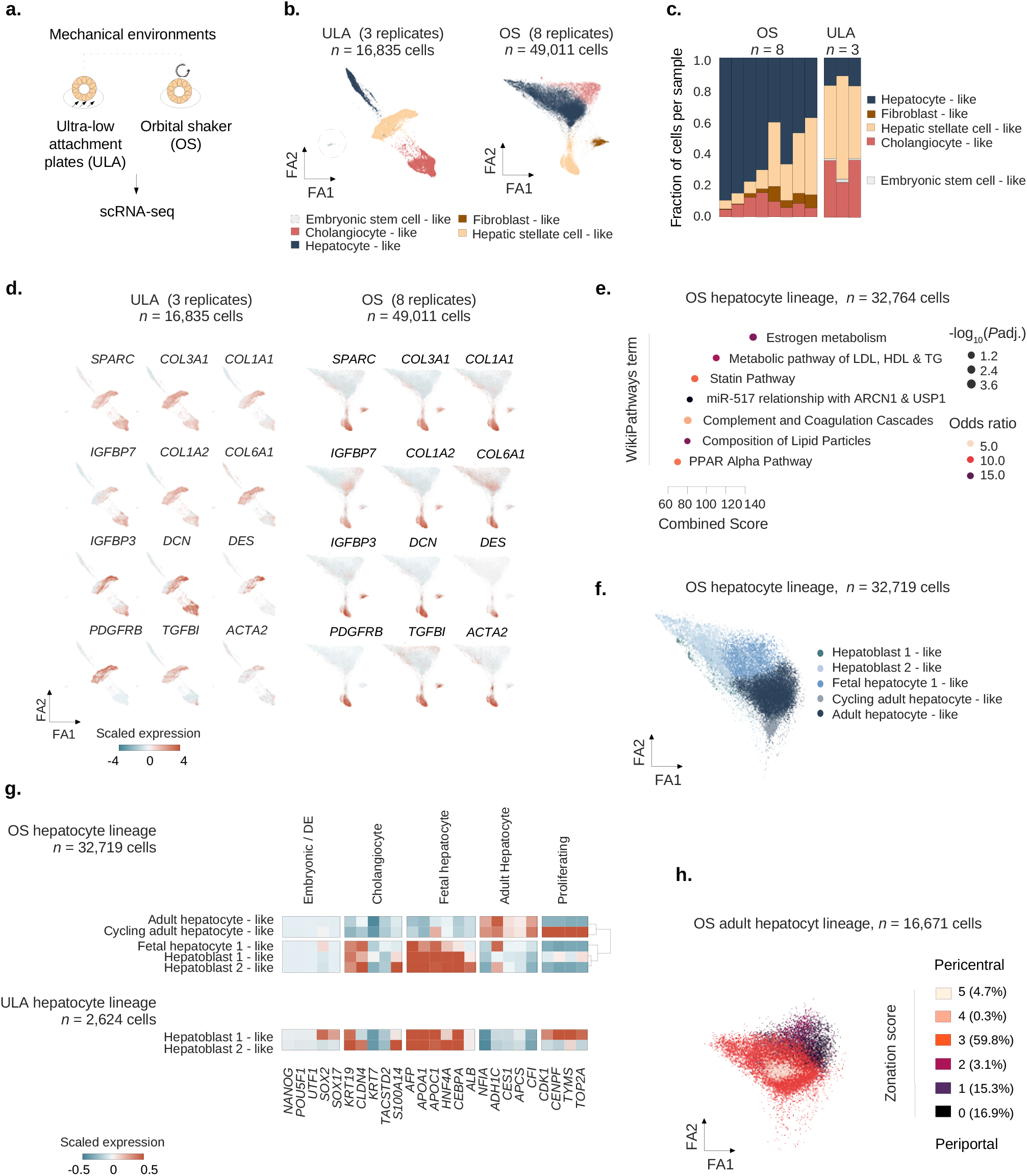
Improved hepatocyte-to-HSC ratio may account for fibrotic and inflammatory responses observed HLOs from different mechanical culture contexts. **a.** Overview of the two mechanical environments selected for 10X scRNA-seq. **b.** ForceAtlas2 plots mapping cells from control ULA-HLOs (cultured in <u>u</u>ltra low <u>a</u>ttachment plates, *n =* 16,835 cells, *n* = 3 replicates) and control OS-HLOs (cultured in <u>o</u>rbital <u>s</u>haker, *n =* 49,011 cells, *n* = 3 replicates). Circled line highlights the embryonic stem cell-like population, which is only present in ULA-HLOs. **c.** OS-HLOs show a relative reduction of HSC- and cholangiocyte-like cells while the fraction of hepatocyte-like cells is increased. Barplots showing the cell cluster proportions as the fraction of total cells per sample for each of the individual OS- and ULA-HLO replicates. The color indicates the cell type as annotated in b., and cell types are listed in the order they are displayed. **d.** The distribution of cells expressing HSC markers is altered in OS-HLOs. ForceAtlas2 representations of the cells from b., colored by the mean scaled expression of HSC marker genes. **e.** Doplot showing the top enriched WikiPathways^82^ terms by combined score. Dot sizes correspond to negative decadic logarithm of the adjusted *p*-value, colors represent the odds ratio. Terms have been shortened for readability. **f.** ForceAtlas2 plots mapping hepatocyte-like cells from control OS-HLOs. Cells were annotated to the human fetal liver development atlas^51^. *n* = 32,719 cells (*n* = 8 controls). **g.** HLOs exposed to the OS-environment show KRT19^low^ hepatocyte subpopulations expressing mature hepatocyte markers. Matrixplot shows the scaled mean expression for marker genes for stages of hepatocyte development for OS-hepatocyte lineage subclusters (top) as annotated in f. in comparison to ULA-hepatocyte-like subpopulations (bottom). Marker genes (bottom) are sorted by cell type (top). Dendrogram representing hierarchical clustering for lineages with > 2 hepatocyte-like subgroups is shown on the right. ULA: *n* = 2,624 cells (*n* = 3 controls), OS: *n* = 32,719 cells (*n* = 8 controls). **h.** Adult hepatocyte-like cells from HLOs exposed to the OS-environment display hepatocyte zonation. ForceAtlas2 plots mapping OS-adult hepatocyte-like cells from f. Color indicates the hepatocyte zonation score as calculated based on marker genes from a human adult liver scRNAseq reference^36^ (Methods). *n* = 16,671 cells (*n* = 8 controls).

We next focused on the hepatocyte lineage and performed pathway enrichment analysis as performed previously for ULA-hepatocyte-like cells, which had displayed enrichment for genes involved in lipid metabolism and glycolysis (**Fig. 1f**). OS-hepatocyte-like cells also showed enrichment in genes involved in specialized hepatocyte metabolic processes, such as estrogen metabolism and coagulation-related pathways (**Fig. 3e**). We then annotated OS-hepatocyte-like cells with recently published marker genes for hepatocyte development^51^ (**Fig. 3f**), which revealed a mix of hepatoblast-, fetal hepatocyte-, and adult hepatocyte-like cells. A cluster mapping close to the adult-hepatocyte-like cells containing cells in G2M phase was termed cycling adult hepatocyte-like cells (Supplementary Fig. 4a). We next evaluated the expression of marker genes for normal hepatocyte development stages^51^ across the jointly preprocessed and normalized cells from OS- and ULA-HLOs (**Fig. 3g**). Unlike OS-HLOs which contained clusters across hepatocyte development, ULA-HLO clusters were all categorized as hepatoblast-like. Consistent with a less mature phenotype, SOX2 and *SOX17* were still expressed in one ULA-HLO hepatocyte cluster. Furthermore a broad proportion of ULA-HLO hepatocytes expressed proliferation markers, in contrast to only two hepatocyte subclusters in OS-HLOs. Most importantly, OS-HLOs displayed a major sub cluster of KRT19^low^ cells expressing adult hepatocyte-markers *NFIA, ADH1C, CES1, APCS* and *CFI*. To understand the zonal composition of the non-cycling adult-hepatocyte-like cells, we sub-clustered the 16,671 adult hepatocyte-like cells and calculated a hepatocyte zonation score based on marker genes from a human adult liver scRNAseq reference^36^ (Methods). This revealed adult hepatocyte-like cells representing periportal, interzonal, and pericental zones (**Fig. 3h**), with a bias toward interzonal hepatocytes. Together, these analyses suggest a more mature transcriptional landscape of hepatocyte-like cells developed with OS culture conditions.

### Oleic acid, palmitic acid, and TGF-β1 induce distinct inflammatory and fibrotic responses at the single cell level

To dissect cell-type-specific transcriptional injury-response patterns, we treated HLOs cultured on an orbital shaker with OA, PA, or TGF-β1 for four days before performing scRNA-seq (**Fig. 4a**). We analyzed 82,467 cells after QC (Methods) from 14 HLO samples, annotating cell types as described previously. We next incorporated hepatocyte sub-stages (Fig. 3g) and sub-clustered cholangiocyte-like cells in order to resolve shifts within major HLO populations under treatment conditions. This yielded two new sub-clusters with expression patterns associated with ductal cell (DC)-like, and smooth muscle cell (SMC)-like cells (**Fig. 4b, left**, Supplementary Fig. 4).

**Fig. 4.**
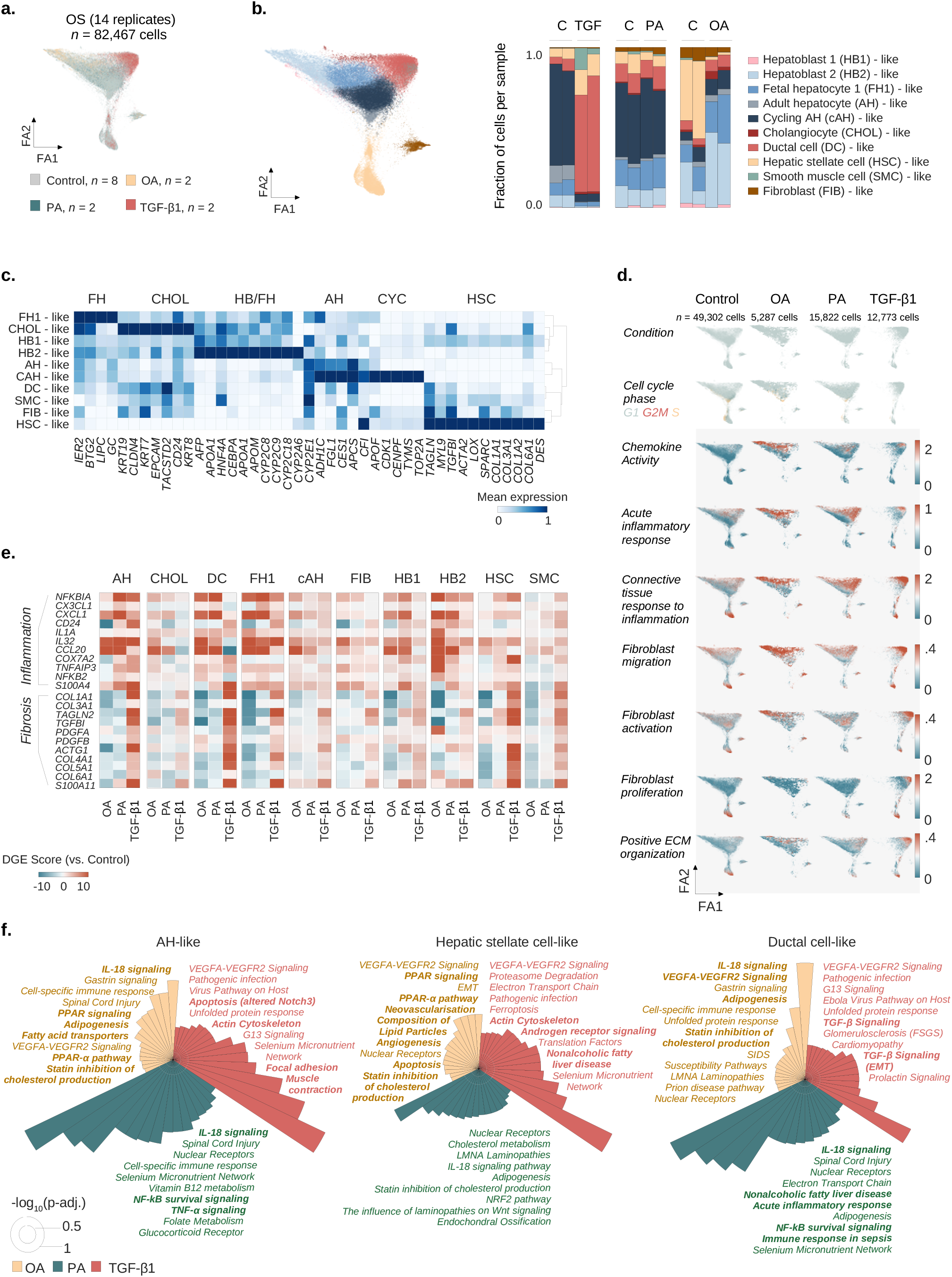
Oleic acid, palmitic acid, and TGF-β1 induce distinct inflammatory and fibrotic responses at the single cell level. **a.** ForceAtals2 representation of cells from OS-HLOs treated for four days with OA (500 µM), PA (500 µM), or TGF-β1 (10 ng/ml), and their respective controls, colored by treatment condition. *N* = 82,467 cells from 14 replicates (*n* = 2 replicates per condition, *n* = 8 controls). **b.** ForceAtals2 representation from a., colored by cell type as annotated with the ScType^40^ database and hepatocyte-like cells resolved to the human fetal liver development atlas^51^ annotation (Methods). Barplot to the right shows cell cluster proportions as the fraction of total cells per sample for each of the individual replicates. Color encodes the cell type annotation, and annotations are listed in the order they are displayed. **c.** Matrixplot shows the scaled mean expression for marker genes in each cluster from b. Canonical marker genes (bottom) are sorted by cell type (top). Hierarchical clustering is represented by the dendrogram on the right. *N* = 82,467 cells from 14 replicates (*n* = 2 replicates per condition, *n* = 8 controls). AH, adult hepatocyte; CYC, cycling; CHOL, cholangiocyte; FH, fetal hepatocyte; HB, hepatoblast; HSC, hepatic stellate cell. **e.** FFAs induce an inflammatory signature, TGF-β1 induces a fibrotic signature, and OA ameliorates the fibrotic signature in HLOs. Cell clusters were separated for expression analysis (top). Differential expression score (pairwise comparison between treatment specific controls and treatments, Methods) for selected genes associated with fibrosis and inflammation for cell types indicated on the top. Differential gene expression (DGE) data are provided in Supplementary Table 5. AH, adult hepatocyte-like; CHOL, cholangiocyte-like; HSC, hepatic stellate cell-like; SMC, smooth muscle cell-like; FIB, fibroblast-like; DC, ductal cell-like; HB1, hepatoblast 1-like; HB2, hepatoblast 2-like; FH1, fetal hepatocyte 1-like. **c.** OA and PA induce inflammatory signals, and TGF-β1 induces expanded fibrotic signatures. ForceAtlas2 plots show GO term^54,83^ scores for selected categories related to inflammation and fibrosis (labeled, left) for HLOs treated with OA, PA, TGF-β1, and controls (labeled, top). Increased expression is indicated by a shift from blue to red. Cell cluster annotations are provided in b. Scale bars indicate the expression score. *N =* 14 replicates (*n* = 2 replicates per condition, *n* = 8 controls). Cell numbers are indicated. **f.** Top enriched WikiPathways^82^ terms for treatment-specific differentially expressed genes in three HLO cell populations corresponding to b., as labeled. Circular bar plots display the negative decadic logarithm of adjusted *p*-values for the respective terms. Cut-off for plots is an adjusted *p*-value below 0.05. White circles indicate negative decadic logarithm of adjusted *p*-values of 0.5 (inner white circle) and 1 (outer white circle). Terms have been shortened for readability, full lists and further cell clusters are provided in Supplementary Fig. 5.

We then evaluated how cell type distributions change with each condition. TGF-β1-treated HLOs displayed an increase in HSC-related populations. HSC-like cells increased from less than 10% in controls (5.4% and 6.6%) to greater than 10% (15.7% and 13.5%) with TGF-β1 treatment. DC-like cells increased from 4% and 7% in the controls to 61% and 72%, potentially mirroring the ductular reaction seen in chronic liver injury. SMC-like cells increased from 0.2% to 14% and 37% with TGF-β1 treatment. In contrast, hepatocyte-like populations decreased by more than 10-fold from 59-63% to ∼5%, and a near-total loss of cycling cells was observed (**Fig. 4b, right**, Supplementary Table 3), in line with previously reported cell cycle arrest through TGF-β1^52,53^. While the analysis shows an expansion of HSC- and ductal-like cells in response to TGF-β1, as observed in the development of liver fibrosis^5,24,25^, no significant alterations in cell population proportions were observed with PA treatment.

In contrast, OA-treated HLOs showed a dramatic reduction of HSC-like cells from 38% and 48% to 1.9% and 1.6%, even though HSCs were comparatively high in this batch of OS- HLOs (see for Fig. 3c). SMC- and fibroblast-like cells were not affected. Conversely, adult-, fetal-, and most strikingly hepatoblast 2-like cells expanded under OA-treatment, the latter from 26% and 8% in controls to 47% and 39% in OA-treated HLOs. This suggests that OA-treatment in HLOs causes a dramatic reduction of HSC-like cells and expands hepatocyte-like cells, specifically hepatoblast-like sub-populations.

Evaluation of the marker gene profiles revealed that DC-like cells were positive for *KRT19* and *HNF4A* as previously reported for this bi-potent cell type induced under conditions of chronic liver injury^25^. Interestingly, DC-like, SMC-like, and fibroblast-like cells acquired a myofibroblast-like gene expression signature (*TAGLN, MYL9, SPARC*) accompanied by a reduction in the expression of ductal markers *EPCAM, CD24* and *CLDN4* (**Fig. 4c**).

To define gene expression signatures activated with each treatment, we evaluated injury-response scores based on gene ontology^54^ term scoring. OA and PA-treated HLOs showed induction of immune response-related scores when compared to their controls and were enriched for chemokine-activity associated genes, as well as fibroblast migration and activation signatures (**Fig. 4d**). TGF-β1 treatment increased scores for fibroblast proliferation and positive regulation of ECM organization along with alterations in cell cycle phase distributions, while these scores remained mostly unchanged in the OA and PA-treated HLOs. Cells with induced inflammatory scores under OA-treatment were primarily hepatoblast-like cells, while PA-treated cells were more broadly enriched for inflammatory scores. Induction of pro-fibrotic scores with TGF-β1 occurred mostly in the mesenchymal clusters (HSC-, fibroblast-like cells) and DC-like cells.

Differential gene expression analysis per cell cluster further displayed injury-specific signatures depending on the respective cell type. The fibrotic module included *COL1A1, TAGLN2,* and *TGFBI* and was induced across TGF-β1 treated cell types, with AH-, DC-, and HSC-like cells showing the strongest relative induction (**Fig. 4e**). The inflammatory module of differential genes included Interleukin 32 *(IL32)* (the top up-regulated gene in NAFLD^55^) and other canonical inflammatory genes including *NFKBIA* and *CCL20* (**Fig. 4e**, Supplementary Table 5), showing the strongest relative induction in AH-, and hepatocyte precursor-like cells treated with PA, followed by DC- and HSC-like populations. Interestingly, OA caused a relative reduction of the fibrotic signature across almost all cell types (indicated by a shift towards blue color). Further, the inflammatory module showed only moderate changes in OA-treated adult hepatocyte-like cells, while the expanding precursors, especially the hepatoblast 2-like cells increased the expression of inflammatory genes.

To contextualize differential gene expression more broadly, we performed cell-type resolved pathway analysis and found TGF-β1-treatment induced apoptosis-, and cytoskeleton-related pathways in AH-like cells, while HSC-like cells were enriched for genes associated with NAFLD (**Fig. 4f**, Supplementary Fig. 5). Strikingly, PA-treated DC-, hepatoblast 1 and 2-, and fibroblast-like cells displayed the NAFLD pathway among their top ranked enriched terms. Additionally, genes associated with TNFɑ-signaling were induced in PA-treated AH-like cells. OA-treatment resulted in lipid-metabolism related hits. Together, these analyses show that PA and TGF-β1 mirror NAFLD-associated inflammatory responses, while the OA-related signature is dominated by genes involved in adipogenesis, fat metabolism and steatosis alongside a general inflammatory response mainly driven by hepatoblast-like cells.

### OA induces an interactome distinct from crosstalk observed with TGF-β1 and PA treatment

To understand the cell-cell interactions in OA, PA and TGF-β1 injury models, we utilized CellPhoneDB^56^ and assessed the relative abundance of interactions between each cell type. Overall, the relative induction of interactions increased from OA-, to PA-, to TGF-β1-treated HLOs (**Fig. 5a**). OA moderately induced interactions between hepatoblast 2-, cholangiocyte- and DC-like cells (HB2, CHOLs, DCs). PA-mediated crosstalk was mainly driven by cholangiocyte-like, hepatocyte progenitor-, fibroblast- and SMC-like cells (CHOLs, DCs, HB1, FIBs, SMCs). Only TGF-β1 yielded a strong activation in cell-cell communication driven by SMC- (most likely representing myofibroblast-like cells) and HSCs-like cells. Plotting the proportion of interactions for each cell type revealed the relative reduction of HSCs- and SMCs-involving interactions in OA-treated HLOs, and their relative induction with TGF-β1 (**Fig. 5b**). Hierarchical clustering of the delta values between control and treatment fractions of each interaction suggested that TGF-β1 induced an interaction pattern more distinct from the ones characterizing HLOs treated with PA and OA (**Fig. 5c**, Supplementary Fig. 6a). These evaluations of the interactome reveal a hierarchical increase of the relative interaction changes induced, in particular with respect to fibrosis-related cell types (HSCs, SMCs), from OA-, to PA, to TGF-β1 treatment.

**Fig. 5.**
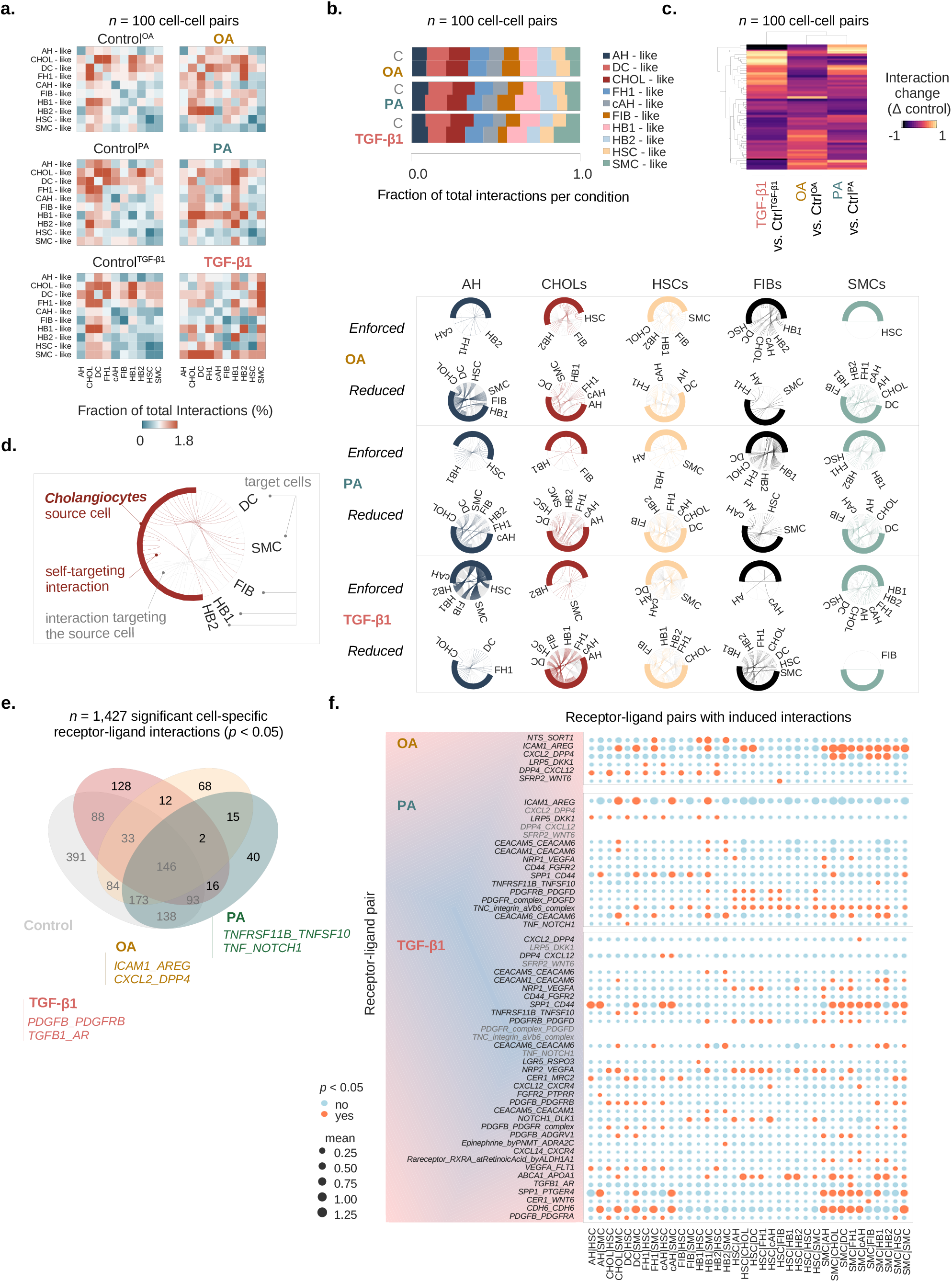
OA induces an interactome distinct from crosstalk observed with TGF-β1 and PA treatment. **a.** CellPhoneDB^56^ analysis of cell-cell interactions in healthy and injured HLOs. Heatmaps show the fraction of total interactions per condition in day 25 OS-cultured HLOs treated with OA, PA, TGF-β1, and their respective controls. AH, adult hepatocyte-like; CHOLs, cholangiocyte-like; DC, ductal cell-like; FH1, fetal hepatocyte 1-like; cAH, cycling adult hepatocyte-like; FIB, fibroblast-like; HB1, hepatoblast 1-like; HB2, hepatoblast 2-like; HSC, hepatic stellate cell-like; SMC, smooth muscle cell-like. *N* = 100 possible cell-cell interactions per condition, *n* = 2 replicates per condition, *n* = 6 controls. **b.** Stacked bar plots show the fraction of total interactions per condition for all cell clusters, where the respective cell is the source of the interaction. Abbreviations are as listed in a. Cell clusters are colored in the order of their appearance. *N* = 100 possible cell-cell interactions, *n* = 2 replicates per condition, *n* = 6 controls. **c.** Clustered heatmap shows the difference in the fraction of total interactions between each treatment and its control group. Dendrograms show the hierarchical clustering by condition (top) and cell-cell interaction (left). *N* = 100 possible cell-cell interactions, *n* = 2 replicates per condition, *n* = 6 controls. **d.** Chord diagrams show the fraction of total interactions per source cell with its respective target cells in HLOs treated with OA, PA, and TGF-β1 relative to their controls. Chord diagrams are divided by interactions that have i) a higher fraction of total interactions in treatment conditions (“enforced”), and ii) interactions with a lower fraction of total interactions (“reduced”) compared to controls. Colored lines indicate interactions initiated by the source cell in each column, while gray lines show interactions of other cell types targeting the source cell. *N* = 100 possible cell-cell interactions, *n* = 2 replicates per condition, *n* = 6 controls. **e.** Venn diagram displays the numbers of intersecting and unique significant (*p*-value < 0.05) cell-type-specific receptor-ligand interactions of HLOs treated with OA, PA, TGF-β1, and controls. Examples for receptor-ligand pair categories containing unique interactions for each treatment are displayed (Methods). *N* = 1,427 significant cell-cell partner-specific receptor-ligand pair interactions, *n* = 2 replicates per condition, *n* = 6 controls. **f.** Dotplots show receptor-ligand pairs with an induced number of significant cell-cell interactions in HLOs treated with OA, PA, TGF-β1, compared to their respective controls. Receptor-ligand pairs with at least two induced interactions are shown. Mean expression (mean of all partners’ individual average expression, Methods) is indicated by the circle size, and significant *p*-values < 0.05 are indicated by orange color. An interaction can be non-significant despite high mean expression among two partnering molecules. Receptor-ligand pairs are shown on the x-axes, and cell clusters are shown on the y-axes. *N* = 2 replicates per condition, *n* = 6 controls.

We next aimed to understand which interactions were enforced or reduced with each treatment when compared its control. We plotted chord diagrams for each cell type and each case, where “enforced” indicates a positive delta value compared to the control and “reduced” indicates a negative delta value (**Fig. 5d**, Supplementary Fig. 6b). This analysis showed that in OA-treated HLOs, particularly SMC-, and HSC-like cells lost more interactions than they gained, and was in line with the general decrease in interaction abundance in these cell types (compare Fig. 4a). In PA-treated HLOs the fraction of SMC-like mediated interactions were shifted in favor of targeting hepatocyte precursors and HSC-like cells. Even though their proportion in cell numbers were not changed with PA (compare Fig. 4b), adult hepatocyte-like sourced interactions were shut down across many interacting partners with PA in favor of enforced communication with hepatoblast 1- and HSC-like cells. In TGF-β1-treatment, we observed an increased fraction of HSC-like and SMC-like cells (Fig. 4b). These cells globally expanded their interactions and shifted their crosstalk specifically towards each other in the presence of TGF-β1. Together, these results suggest OA-mediated reduction in HSC- and SMC-like cellular crosstalk while TGF-β1 promotes crosstalk between these cell types.

As a next step, we identified cell-type resolved receptor-ligand pairs exclusive for each injury type. With PA treatment, receptor-ligand expression involving *TNF superfamily (TNFS)* members was observed, reflecting the induction of previously described molecular mediators of cellular responses to injury and inflammation in the liver^4^. In TGF-β1-treated HLOs, receptor-ligand expression was observed involving *PDGFB* and *PDGFRB,* recapitulating an important hallmark of liver fibrosis *in vivo*^22,57,58^ (**Fig. 5e**, Supplementary Table 6). We further identified *CXL2* and *DPP4* ligand-receptor pair expression signatures indicative of OA-treatment. Inhibitors of the exopeptidase dipeptidyl-peptidase 4 (DPP4) are anti-diabetes drugs and have been shown to reduce steatosis in NAFLD patients^59^, though failing to consistently ameliorate liver inflammation and fibrosis in larger meta analyses^60^. The induction of predominantly NAFL/steatosis-related interacting transcripts is in line with our previous analyses where OA induced steatosis in HLOs (Fig. 2) without fibrosis and had more localized effects on the broad inflammatory response. This evaluation of the interactome reveals groups of receptor-ligand pairs changing in response to inflammatory and fibrotic injury in HLOs overlapping with molecular mediators previously reported *in vivo*.

We then examined receptor-ligand pair expression to explore injury-specific interactions (**Fig. 5f**). These analyses revealed new *DPP4-CXCL12* pairs with OA treatment potentially allowing hepatocyte-like lineages and fibroblast-like cells to cleave CXCL12 expressed by HSC-like cells. The expression of TNF-Related Apoptosis Inducing Ligand (TRAIL) encoded by TNF superfamily member *TNFS10* in HB-like cells in linkage with its decoy receptor Osteoprotegerin (OPG)^61^, encoded by *TNFRSF11B*, in SMC-like cells was exclusively observed in the PA condition, whereas this interaction expanded shifting towards matured hepatocyte-like stages and SMC-like cells with TGF-β1. OPG expression has been suggested as an evasion mechanism from TRAIL-induced apoptosis acquired by activated myofibroblasts in liver fibrosis^62^, and its expansion in loco typico (SMC-like cells) suggests the gradual acquisition of fibrogenic activation in these myofibroblast-related cells. Osteopontin (encoded by *SPP1*) can bind CD44 to regulate cell adhesion and migration^63^. SMC-like cells with PA and TGF-β1 treatment induced autocrine interactions through *SPP1* and *CD44,* consistent with the behavior of activated HSCs in mouse and human NASH livers^64^. With TGF-β1, the SMC-like cell-sourced *SPP1/CD44* interaction further expanded within the mesenchymal targeting HSC-like cells. TGF-β1 treatment also broadly induced *PDGFB/PDGFR* interactions, indicating the acquired potential for hepatocyte-like lineages to signal to HSC-like cells through PDGFRA and hepatocyte- and DC-like lineages to signal to HSC- and SMC-like cells through PDGFRB, in line with the critical role of PDGFB-mediated HSC-activation in liver fibrogenesis^57,58^. We also find less studied targets, including TGF-β1-mediated *TGFB1/Androgen receptor (AR)* interactions. Together, these findings highlight the alignment of subsequent cellular crosstalk changes from OA, to PA, to TGF-β1-treated HLOs to pathways involved in human NAFLD progression. Our analyses further highlight the androgen-receptor pathway as a potential molecular target in fatty acid-induced liver injury and fibrosis.

### Trajectory inference reconstructs the emergence of major HLO lineages and injury-specific terminal states

Progression from steatosis to steatohepatitis and then fibrosis can be understood as a sequential process that includes the emergence of myofibroblast lineages^4,5^. We investigated the projection of our steatosis (OA), steatohepatitis (PA), and fibrosis model (TGF-β1) on a force-directed layout in conjunction with Palantir^65^ trajectory inference analysis to dissect this process.

We first analyzed the hepatic progenitor lineage (**Fig. 6a**) choosing the *ALB*^highest^/*CEBPA*^pos^ hepatoblast-like cell as the early cell (**Fig. 6b, top**). As expected from the immature gene expression signatures in hepatoblast-like cells, pseudotime increased towards the adult hepatocyte-like cluster (**Fig. 6b, top**). Three terminal states were identified, including the cycling adult-hepatocyte-like state and two DC-like terminal states emerging through the cholangiocyte-like cells (**Fig. 6b, top**). We next identified cells with high differentiation potential (**Fig. 6b, bottom**), showing enhanced activity in three defined regions, including one at the hepatoblast/cholangiocyte (arrowhead 1) and two at the cholangiocyte-ductal (arrowhead 2 and 3) interface. Terminal states are defined by a single cell and do not allow us to evaluate the population of cells most closely related to the terminal state. To understand the contribution of each treatment to terminal states, we selected the top 100 terminal cells for each of the three terminal states identified and projected these cells on the force-directed layout (**Fig 6c**, top). Analysis of the cell proportions revealed that the AH-like cycling terminal state (cAH) was primarily composed of cells from the control conditions, while the DC-like states were dominated by TGF-β1-treated cells (**Fig. 6c**, bottom). Performing a similar analysis on the HSC-lineage (**Fig. 6d**), choosing the *IGFBP*^highest^/*DDIT4*^pos^ cell as the early cell, revealed five terminal states in the regions of three HSC-like clusters (referred to as HSC1-3), the fibroblast-like cluster (FIB), and one SMC-like cluster (**Fig. 6e, top**). The three regions of high differentiation potential corresponded to the interfaces between these major mesenchymal cell types (**Fig. 6e, bottom**, arrowheads 1-3). Analysis of the cell proportions revealed that the FIB and HSC2 terminal states were mostly represented by cells from the control condition, while the HSC1 and HSC3 were primarily composed of cells from control and TGF-β1-treated cells. In contrast, SMC terminal state was largely represented by TGF-β1-treated cells (**Fig. 6f, bottom**). Together, these analyses suggest that only TGF-β1-treatment favored the emergence of a myofibroblast-like terminal state, consistent with a canonical feature of fibrosis pathology^5^.

**Fig. 6.**
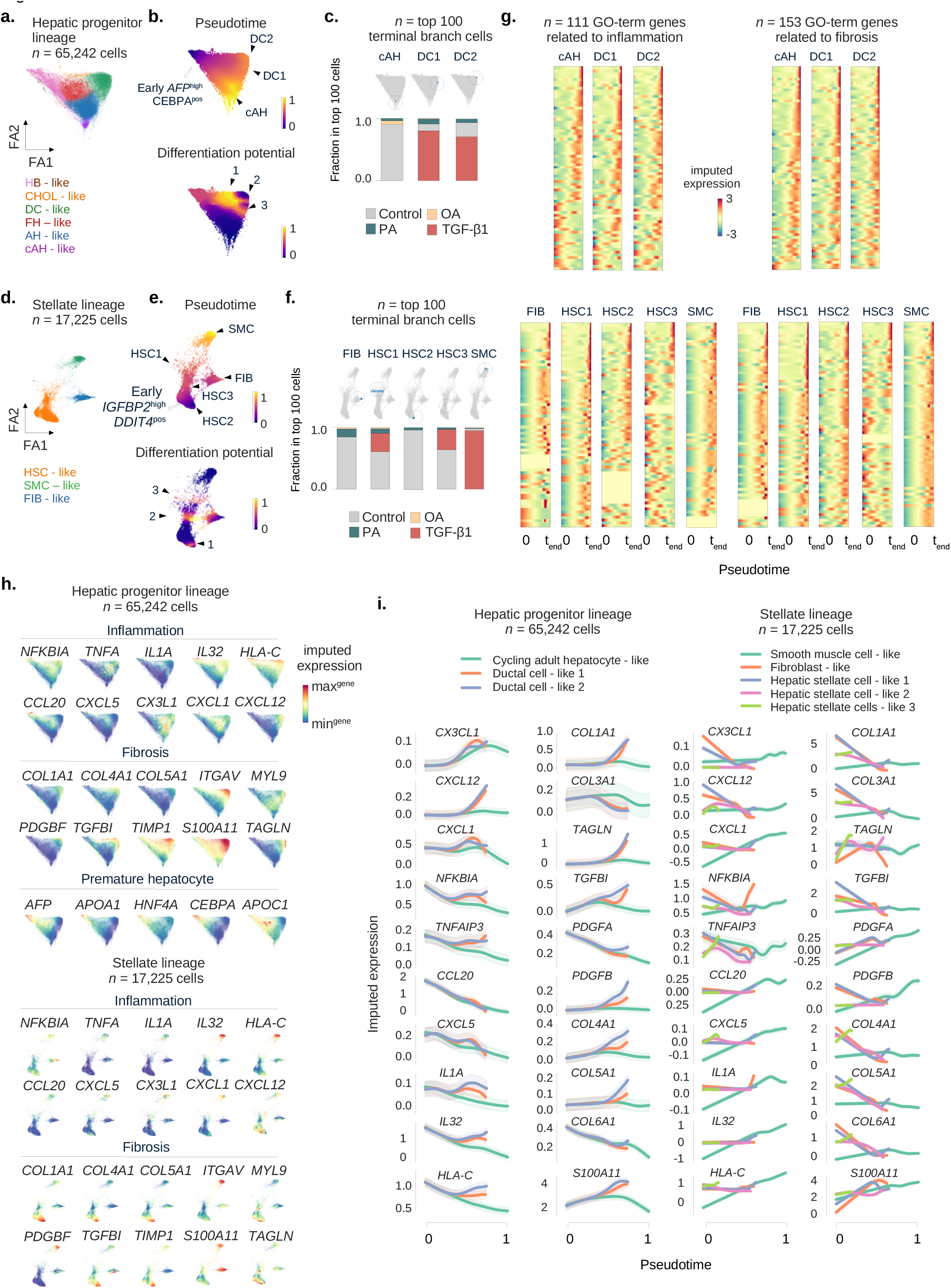
Trajectory inference reconstructs the emergence of major HLO lineages and injury-specific terminal states. **a.** ForceAtlas2 representation of the hepatic progenitor lineage composed of cells from healthy controls and OS-HLOs treated with OA, PA, and TGF-β1, colored by cell type. *N* = 65,242 cells from 14 replicates (*n* = 2 replicates per condition, *n* = 8 controls). **b.** ForceAtlas2 representation from a., colored by Palantir trajectory inference results based on the ForceAtlas2 representation, showing pseudotime (top) and differentiation potential (bottom). For pseudotime, black arrowheads indicate three pseudotime maxima, and the white arrowhead identifies the early cell (selected based on the highest level of *AFP* expression among cells expressing *CEBPA,* both of the genes marking hepatocyte precursors^51^). Regions of high differentiation potential are indicated by arrowheads 1-3. **c.** Projection of the top 100 terminal branch cells (sorted by branch probabilities) for each terminal state identified by Palantir on the ForceAtlas2 representation from a. (top). Corresponding barplots below displaying the relative distribution of treatment conditions among the top 100 terminal branch cells for each terminal state. **d.**, **e.**, and **f.** display the plots from a-c for the HSC lineage, in this case the early cell is selected based on the highest level of *IGFBP2* expression among cells expressing *DDIT4*, as a marker of fetal HSCs^51^. *N* = 17,225 cells from 14 replicates (*n* = 2 replicates per condition, *n* = 8 controls). **g.** Gene expression trends reveal the increase of inflammatory and pro-fibrotic transcripts along the pseudotime ordering towards PA and TGF-β1 dominated terminal states. Heatmaps showing the imputed gene expression over Palantir pseudotime for GO-term^54,83^ derived inflammation (left) and fibrosis (right) related genes sorted by their imputed expression level at each terminal state (indicated on top of each heatmap). Imputed expression levels are indicated by color, y-axes correspond to genes, and x-axes display the pseudotime. Ranked gene lists are provided in Supplementary Table 7. **h.** ForceAtlas2 representations for hepatic progenitor and HSC lineages from a. and d., showing the MAGIC-imputed gene expression of selected inflammation- and fibrosis-related genes from g. enriching towards specific terminal states. Fetal hepatocyte-related transcripts enrich along the trajectory towards the OA-treated hepatoblast-like cells. Projection of the imputed gene expression of transcripts related to inflammation, fibrosis, and the developmental stage of hepatoblasts is indicated by section headings. Color represents the range of imputed expression values for each gene. **i.** Gene expression trends for hepatic progenitor (left three columns) and HSC (two right columns) lineages of representative enriched genes from g. Pseudotime is indicated on the x-axes and normalized expression is shown on the y-axes. Cell types are indicated by color (top) and can be identified in a. and d. Note that not not all terminal cells reach the maximum pseudotime value of 1 and therefore terminate their trend line before the x-axis maximum (compare b. and e.).

To examine the differentiating regions as potential representations of healthy-to-injury transitions, we next applied MAGIC^66^ to generate imputed data and visualize the expression of relevant genes along the differentiation trajectory with a focus on the acquisition of “pro-fibrotic” and “pro-inflammatory” gene expression signatures. We selected genes based on GO-terms^54^ and sorted the genes according to their imputed expression at the final pseudotime to visualize whether these signatures were enriched towards each terminal state (**Fig. 6g**, Supplementary Table 7). For the hepatic progenitor lineage, this analysis revealed an increase of pro-fibrotic gene signature enrichment across pseudotime for cAH- and DC-like terminal states (**Fig. 6g, top**). In the HSC-like lineage, the SMC-like terminal state enriched a broad spectrum of pro-inflammatory and pro-fibrotic gene signatures along pseudotime, with variable enrichment for HSC-like and fibroblast-like cells (**Fig. 6g, bottom**). Interestingly, OA-treated cells were hardly represented across the cells constituting the extrema of trajectories leading to terminal states with fibrotic signatures.

To resolve transcripts specific to each terminal state, we projected the imputed expression of genes enriched in the above analyses or related to hepatocyte precursor states on the force directed layout for each lineage (**Fig. 6h**). In the hepatic progenitor lineage, inflammatory transcripts *NFKBIA*, *CXCL6* and *CXCL1* increased towards the TGF-β1-dominated DC-like terminal state. We observed a targeted diffusion distribution of pro-fibrotic genes towards the same cells, again corresponding to the TGF-β1-treatment enriched trajectory (e.g., *TGFBI* and *TIMP1*). The hepatoblast-like terminal state (that expanded under OA treatment, Fig. 4b) showed enrichment for hepatocyte precursor defining transcripts *AFP, APOA1, APOC1, HNF4A* and *CEBPA*, providing further evidence that OA favors the expansion of hepatocyte precursor states. Profiling of the HSC-related lineages revealed *IL32* and *HLA-C* as components of the inflammatory, and *ITGAV* and *S100A11* as components of the fibrotic signature towards the SMC-like terminal state, which was overrepresented by TGF-β1 treated cells. These transcripts are also characteristic of activated HSCs and a hallmark of fibrosis *in vivo*^55,67^, suggesting the emerging SMC terminal state represents the activation of these HSC-like cells.

To better understand gene expression signatures accounting for deviating terminal states observed in injured HLOs, we next projected lineage-specific gene expression trends over pseudotime (**Fig. 6i**). This highlighted pro-fibrotic and pro-inflammatory genes characterizing individual terminal states of the hepatic progenitor lineage, which were the cAH- and DC-like state (**Fig. 6i**, two left columns; green, orange and blue lines, respectively). In the HSC-related lineages (**Fig. 6i**, two right columns), inflammatory signatures accompanied the trajectory towards the TGF-β1-specific SMC-like terminal state (green line), along with fibrosis-related genes such as *PDGFB, S100A11* and *TAGLN*. Pseudotime is indicated on the x-axis, with a value of 1 representing the greatest pseudotime. Some cell populations terminate before 1 based on pseudotime calculations for each terminal cell state. Together, these results support the presence of inflammatory and fibrotic signatures on the path towards fibrosis being modeled by PA and TGF-β1 treatments, and underscore the induction of premature hepatocyte stages by OA.

### NAFLD severity prediction gene signature scores progressively increase from OA, to PA, to TGF-β1

We next investigated how changes in gene expression following treatment of HLOs with OA, PA, and TGF-β1 relate to the development of fibrosis in patients with NAFLD. We applied the 26 and 98 gene signatures established to predict fibrosis in NAFLD^68^ to bulk gene expression in HLOs. This analysis revealed a significant increase in the score for both gene signatures with all treatments, with the mean score increasing to the greatest extent with TGF-β1 treatment, followed by PA-, and OA-treatments (**Fig. 7a**). We then plotted expression signatures for individual genes that demonstrated the greatest dynamic range in expression across clusters (**Fig. 7b**). Many of these genes show a trend of induction from control, to OA, to PA, to TGF-β1, including *TAGLN2, CXCL6*, *CYTOR*, and *S100A11*, while other genes, such as *S100A4* show the highest expression with OA treatment. These results demonstrate a step-wise induction of NAFLD disease progression scores across treatment conditions in the HLO system.

**Fig. 7.**
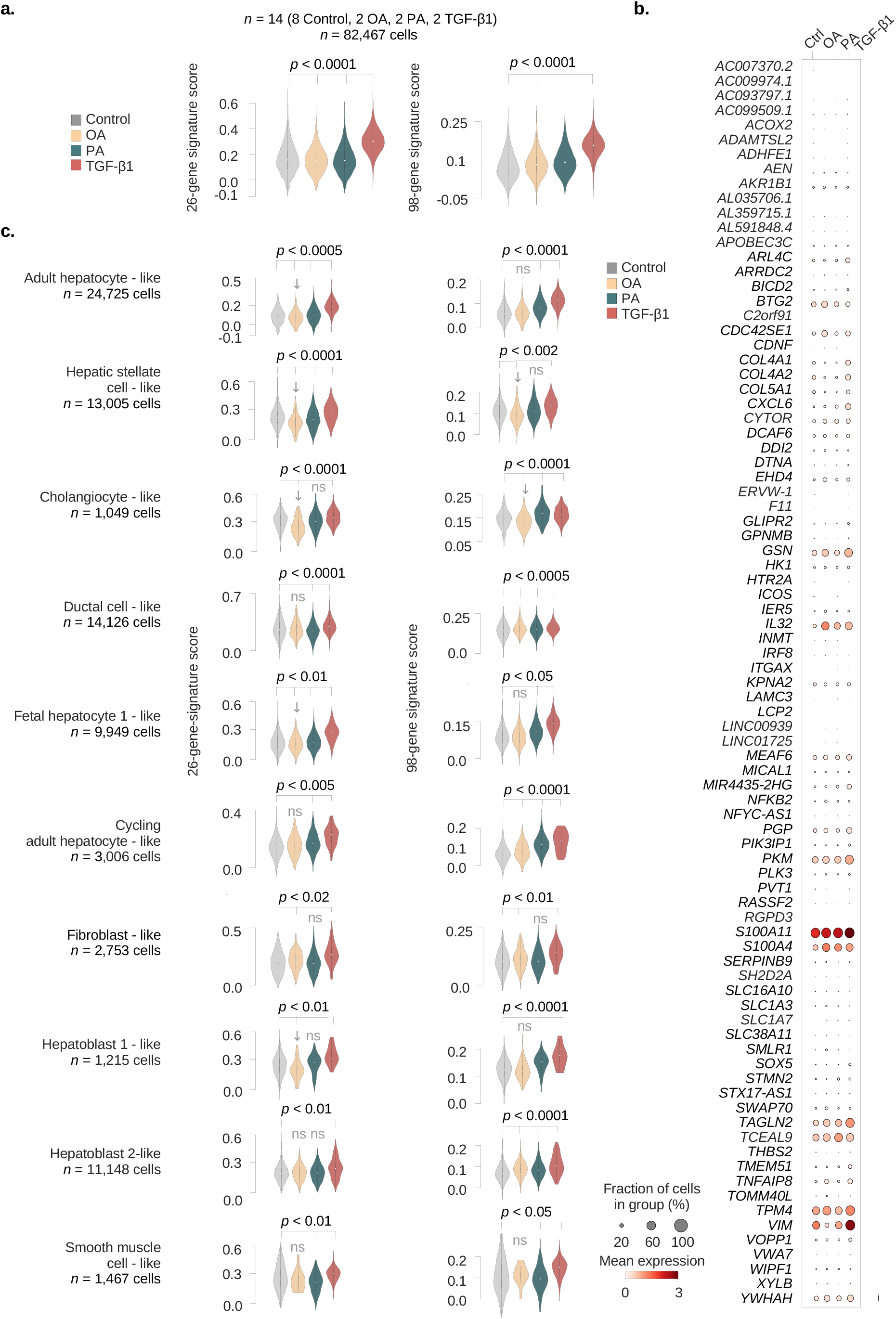
NAFLD severity prediction gene signature scores progressively increase from OA, to PA, to TGF-β1. **a.** Application of 26 and 98 gene signatures^68^ to predict fibrosis stages across NAFLD to control and injured OS-HLOs. *P*-values derived from Kruskal-Wallis test (two-tailed) with post hoc Conover’s test followed by Bonferroni correction. Ns, non-significant. *N* = 82,467 cells from 14 replicates (*n* = 2 replicates per condition, *n* = 8 controls). **b.** Dotplot shows the scaled mean expression of commonly expressed genes from the 98-signature score across treatment conditions. Dot sizes correspond to the percentage of cells expressing the respective gene in each condition, and the level of expression is indicated by color. **c.** Deconvolution of the scores from a. to individual cell types as annotated in Fig. 4b. *P*-values derived from Kruskal-Wallis test (two-tailed) followed by a post hoc Conover’s test with Bonferroni correction are indicated. *N* numbers are indicated below cell cluster names. Ns, non-significant.

We then evaluated the 26 and 98 gene signatures at the level of cell types. The 26 and 98 gene signatures increased across the major cell types of adult hepatocyte-, HSC- and cholangiocyte-like cells with significance in at least one of the two signature scores for PA and TGF-β1 treatment (**Fig. 7c**). In contrast, we observed a reduction in both 26 and 98 gene signatures with OA treatment for HSC- and cholangiocyte-like cells and a decrease in the 26 gene signature for adult hepatocyte-like cells. The main contributors to the OA-related NAFLD signature induction were hepatoblast 2-like cells. To further dissect the contribution of individual cell types to the score enrichment, we next plotted the relative expression of individual genes per cell cluster across treatment conditions (Supplementary Fig. 7). Adult hepatocyte-like cells in HLOs treated with TGF-β1 showed higher expression of *S100A11*, while OA and PA induced *TCEAL9* in this population. HSC-like cells showed induction of *COL4A1*, *COL4A2, COL5A1, PKM, VIM,* and *TPM4* with TGF-β1 treatment, while OA and PA treatment resulted in increased expression of the inflammatory signature gene *IL32* in both epithelial and mesenchymal clusters. Together, these results indicate the gradual acquisition of gene signatures linked to disease progression in NAFLD across the major HLO cell types with PA and TGF-β1, additionally resolving a mixed response in the OA condition, where hepatocyte precursors were the main drivers of gene expression changes observed with NAFLD progression, while AH- and HSC-like populations lost the NAFLD fibrosis signature.

## Discussion

While studies have begun to decipher the contribution of individual cell types to the development of cirrhosis in humans^22^, our understanding of gene expression programs and the interaction between cell types during evolving liver fibrosis in the context of NAFLD is limited. Conventional 2D cultures fail to provide cellular complexity, and animal models come with ethical constraints and do not involve human cells. Recent multi-lineage *in vitro* organoid systems, such as human liver organoids (HLOs), hold the promise to bridge the gap between cell culture and clinical observations^69^. In HLOs, NAFLD and fibrosis have been studied so far by oleic acid (OA) treatment^27^ and genetic manipulation^28^. However, saturated FFAs, like 16:0 palmitic acid (PA), show a tighter association with NAFLD^70^ and more efficiently induce hepatocyte damage^71,72^. Despite the critical importance to choose the most suitable HLO-NAFLD model, a systematic evaluation of the efficacy on NAFLD transcriptome induction among lipotoxic agents in HLOs has not yet been performed.

Here, we examined the effect of OA-, PA-, and TGF-β1-mediated injury in HLOs and the ability of each condition to model the progression of steatohepatitis and fibrosis caused by NAFLD-induced liver disease. This analysis covered histologic, phenotypic, and gene expression studies, including the creation of a reference of ∼100,000 single-cell transcriptomes of injured and healthy NAFLD-HLOs. We find that OA induces patterns of gene expression that are reflective of benign steatosis, while the inflammatory changes of steatohepatitis are induced with PA, and the combination of steatohepatitis and fibrosis is most prominent with TGF-β1.

While OA induced steatosis, inflammatory changes were only observed in hepatocyte precursors. Unexpectedly, OA treatment not only failed to induce *COL1A1* mRNA measured by qPCR and reduced intra-organoidal collagen levels as quantified by Sirius red stainings, but it also significantly ameliorated fibrotic gene signatures across cell types and largely reduced the HSC population in our scRNAseq data. In contrast, PA induced a more robust steatohepatitis signature from more mature hepatocyte sub-populations, while having minimal effect on HSC population size or gene expression. TGF-β1 was the only agent tested that generated a global fibrotic phenotype including collagen type 1 induction. Additionally, only TGF-β1 treatment resulted in the expansion of the *KRT19/HNF4A*+ DC-like population, consistent with the reaction observed with chronic liver injury *in vivo*^23–26^. It is not clear why previous studies in HLOs^27^ show a more marked fibrotic change with OA treatment, but it may reflect differences in culture conditions and/or the presence of a small number of Kupffer-like cells, which we do not observe in our HLOs. However, the conclusion that PA is a more effective model for steatohepatitis in HLOs is consistent with both human and mouse studies.

OA, a monounsaturated fatty acid, is the major component of olive oil^73^. Olive oil is an integral component of the Mediterranean diet (MD)^74^, and growing evidence over the past decades has demonstrated health benefits from the MD, attributed to its high olive oil content^74^. The beneficial effects on cardiovascular disease are well established^75,76^, and a recent meta-analysis of randomized controlled trials highlights the positive effects of the MD in NAFLD, with the observation of reduced ALT levels and liver stiffness^77^. *In vivo* studies in mice and rats have also demonstrated that OA protects against inflammatory and fibrotic effects in models of NASH^14,15,78^. The observation that OA reduces the fraction of HSCs and induces an almost pan-cellular reduction of fibrotic signatures in HLOs, could help explain the beneficial effects observed in these studies.

Differential effects have been observed for OA and PA *in vitro* that also indicate OA could reduce fibrosis. While OA and PA induce steatosis in hepatocytes, and OA is observed to have a more pronounced effect on steatosis than PA in some studies and similar effects in other studies^12^, PA more potently induces cytotoxicity and apoptosis compared to OA^71,72^. Additionally, evidence for OA-mediated protection from lipotoxicity in cultured hepatocytes has been observed^71^. Treatment of HSCs directly with OA leads to reduced collagen, ɑSMA, and vimentin levels and inhibits proliferation^79,80^. In contrast, PA has been linked more directly to fibrosis, as conditioned media from PA-treated hepatocytes induces fibrotic gene expression in HSCs^81^.

Since the involvement of immune cells is an important component in NAFLD development, this constitutes a limitation of current HLO model systems. Further studies will be necessary to i) develop HLO models of greater cell type variety and maturity and ii) continuously evaluate their capacity to model human pathologies.

In summary, we herein provide a systematic approach to benchmark NAFLD-HLOs, including single cell analysis, which will serve as a reference atlas for future studies. The interplay between inflamed, steatotic hepatocytes, cholangiocytes, and HSCs provided in the HLOs demonstrate the antifibrotic effects of OA observed *in vivo*, and suggest they are driven at least in part by a reduction of HSCs. This may be an underexplored mechanism by which OA exerts beneficial effects in animal models and NAFLD patients. Our results also provide evidence for the rationale to favor PA over OA to model steatohepatitis in HLOs and to use TGF-β1 to study the combination of steatohepatitis and fibrosis observed in later stage disease.

## Methods

### hPSC and HLO culture

H1 hPSC cells (WA01) were obtained from WiCell. ESCRO approval was received from Massachusetts General Hospital. hPSCs were maintained in mTeSR medium (StemCell Technologies) on Matrigel (Corning, 354230) as previously described^84^ and split with Accutase (Thermo Fisher Scientific) upon 80-90% confluence. For passage and thawing, Rho-associated kinase (ROCK)-Inhibitor (StemCell Technologies, Y-27632) was added to mTeSR at 10 µM. For the differentiation of human liver organoids (HLOs), cells were split at a ratio of 1:10 and cultured until ∼85% confluent before HLOs were differentiated as previously described^27,34^. On day 21, HLOs were either kept in Matrigel domes or isolated. To perform isolation, HLOs were incubated in DPBS(-/-) on ice for 15 minutes followed by repeated manual dissociation with a P1000 at 4°C followed by centrifugation at 190 x *g* for three minutes, removal of supernatant, and resuspension in 10 µL final medium per receiving well of a 24-well-plate. Isolated HLOs in media were either plated to i) 1% agarose coated plates ii) plates on an orbital shaker at 80 revolutions per minute, or iii) hydrogel plastics ultra low attachment surface 24-well plates (Corning, 3473).

### Liver injury induction

Injury media solutions based on Hepatocyte Culture Medium (HCM, Lonza, CC-3198) were prepared to obtain solutions of TGF-β1 (10 ng/mL and 25 ng/mL, R&D Systems 240-B-002), palmitic acid (PA, 500 µM, Sigma Aldrich P0500), and oleic acid (OA, 400 and 800 µM, Sigma Aldrich O1383). HLO media was changed to injury media on day 21. Isolated HLOs were resuspended in the final injury media at the final step of the isolation process (see above) and kept in solution for four days and harvested on day 25. Controls for TGF-β1 were cultured in HCM. PA was dissolved into HCM with 10% BSA and 1% ethanol before dilution to final concentration in HCM. OA was dissolved into DPBS(-/-) with 12.5 mM NaOH and 1.67% BSA at 8 mM before dilution to final concentration in HCM. PA and OA controls were prepared accordingly, omitting the initial step of dissolving the active agent in the carrier solutions.

### Contraction assay

Day 20-21 HLOs in 50 µL Matrigel domes cultured in HCM were exposed to TGF-β1 at 10 ng/mL and 25 ng/mL. Images of plates with scale bars were taken prior to treatments and at day five of incubation. Measurements were performed with ImageJ (version 1.53a) by manually selecting the HLO/matrix drop areas. All distances in mm² were normalized to the mean of the control areas in mm², and two-tailed Kruskal-Wallis-statistics were performed on all groups with a Conover post hoc test.

### qPCR

Total RNA was isolated from HLOs to perform qPCR. Briefly, 1 mL medium was removed from 6 well plates. Matrigel domes containing HLOs were carefully detached from the wells with a cell scraper and transferred to a 1.5 mL tube including medium. If isolated HLOs served as starting material, the detachment step was not needed. Matrigel-embedded HLOs were left at 4°C for 15 minutes to dissolve the Matrigel. Then, HLOs were centrifuged for three min at ∼190 x *g* at 4°C (Matrigel-embedded HLOs) or at RT (non-matrigel HLOs or 2D cell layers). Supernatant was discarded, and the pellet was resuspended in 300 µL Trizol (Invitrogen, 15596026), rigorously vortexed or mechanically processed until fully dissociated, and finally incubated for 10 minutes at room temperature and then stored at -80°C. RNA was prepared via phenol-chloroform extraction. For cDNA synthesis, 200 or 500 ng RNA were reverse-transcribed using the iScript gDNA Clear cDNA Synthesis Kit (Biorad, 1725034), and no-RT-controls and mastermix controls were prepared. qPCR reactions were prepared from cDNA at 1:5 dilution with SYBR Green iTaq Universal SYBR Green Supermix (Biorad, 1725120) and qPCR primers at 10 µM in a total volume of 10 µL in a ≥ 40-cycle and melt curve reaction cycler (Biorad, Base #CT009383, Optical Head #786BR2648). All biological samples were measured in technical triplicates.

### qPCR data analysis

All gene Ct means were calculated from three technical replicates. Only samples with housekeeping Ct values below 35 were considered for analysis. Mean Ct values for housekeeping genes β-Actin (*ACTB*) or glyceraldehyde-3-phosphate dehydrogenase (*GAPDH*) were subtracted from target gene means to generate the ΔCt value. The control ΔCt value average was calculated from three biological replicates and subtracted from all experimental ΔCts to render ΔΔCt values. The log2 fold-change was calculated as 2^⁻ΔΔCt^.

### Primer design and validation

Primers were designed using PrimerBlast. Briefly, transcripts of interest were selected by NCBI Reference Sequence ID and the ‘pick primers’ hyperlink function was used to import the cDNA sequence into the PrimerBlast mask. The following settings varied from the defaults: Exon junction span → primer must span an exon-exon junction, PCR-product size → 50-200, allowing primers to bind to variant transcripts of the same gene. Blast was performed against both, RefSeq mRNA (Homo sapiens) and RefSeq representative genomes (Homo sapiens) in order to obtain primers specific to transcript sequence and mRNA rather than gDNA. Primers with i) no additional match in RefSeq mRNA or at least > 4 mismatches and ii) no additional genomic hits or hits with at least > 800 bp product size, were selected for further validation. Next, primers were tested on human liver tissue RNA at a final concentration of 10 µM as described above, and 2% agarose gels of the primer product were prepared with a 100 bp ladder. Primers were selected for further experiments if they showed a single band in the expected size range. The band was cut, cDNA was extracted from agarose and sent for Sanger sequencing, and the product identity was assured by re-blasting. A primer was selected for experiments when the sequence matched the selected sequence of the initially selected transcripts (NCBI BLAST). Primer sequences are provided in Supplementary Table 8.

### Histology and immunohistochemistry

Medium was removed from day 16, day 21 or day 25-26 HLOs treated for four days with pro-fibrotic substances, washed twice with warm DPBS(-/-) and fixed overnight at 4°C with 4% PFA in DPBS(-/-). PFA was removed and the HLOs were washed 2x with DPBS(-/-) before being transferred to 70% ethanol for storage and paraffin embedding. H&E and Sirius red staining was performed on cut and deparaffinized HLOs in line with standard protocols. Anti-human-CEBPα (Sigma Aldrich, HPA052734) and the secondary antibody (HPR anti-mouse) were used at a dilution of 1:200.

### Organoid Sirius red quantification pipeline

The pipeline for HLO-specific Sirius red staining analysis is written in python (≥ version 3), ImageJ (version 1.53a) macro and bash and will be made publicly available upon release at https://github.com/anjahess/sirius_red. First, full slides were cleared from black artifacts. Then, for whole slide quantification, all image areas containing tissue were selected and saved for quantification. The Sirius red quantification module was adopted from the NIH-Image J macro “Quantifying Stained Liver Tissue“ (available at https://imagej.nih.gov/ij/docs/examples/stained-sections/index.html, requested 2020-11-14). Briefly, RGB stacks were generated from images, and the tissue containing area was defined and measured by the *setAutoThreshold(“Default stack”)* function. The Sirius red staining threshold reached from minimum to maximum as retrieved from the *setAutoThreshold* and *getThreshold(min, max)* functions measured in the green channel of the RGB stack, and divided by an experiment-specific brightness factor (0.95 or 1.3, similar across all control and treatment conditions). For single HLO analysis, 200 x 200 µm selection squares were placed around HLOs in the interactive, user-supervised mode and resulting images were processed as described for whole slide scans. Finally, csv-formatted result tables were called from python (version 3.7) and the Sirius red stained area was calculated either per total area or per tissue (measures as the auto-thresholded area in the blue channel). For vacuole quantification, the fraction of stain-free tissue was compared. If normalization was performed, values were calculated as the percentage of mean of all control samples of the respective experiment.

### Single-cell RNA sequencing

For day 21 ULA HLO single-cell RNA sequencing, HLOs underwent a passage step at day 16 and remained in HCM medium with 10% Matrigel on ultra-low attachment plates as previously described^34^. At day 21 one well of a six well plate of free-floating HLOs was collected, washed 1X with warm DPBS(-/-) and briefly dissociated by incubation with 300 µL 0.05-0.25% Trypsin-EDTA for 10 minutes at 37°C (Corning, Ref. 25-052-CI RT). Day 25 orbital shaker-cultured HLOs underwent an additional DBPS (-/-) wash and 10 minutes of trypsinization to allow complete HLO digestion. For each replicate, full single cell dissociation was confirmed manually by light microscopy. After a final DPBS(-/-) wash, HLOs were resuspended in DPBS(-/-), counted after Trypan blue staining with the TC20™ Automated Cell Counter according to a previously determined optimal gating of 7-20 µm and transferred on ice. Library preparation was performed on biological replicates with the highest viability counts. Libraries were prepared according to the 10X ChromiumSingle Cell 3ʹReagent Kits v3 instructions. The library was sequenced by paired-end sequencing on an Illumina® NextSeq 2000 P3 flow cell or the NovaSeq 6000 system according to the manufacturer’s recommendations.

### Single-cell RNA sequencing analysis

*<u>Alignment and quality controls (QCs).</u>* Raw bcl files were converted to fastq by the command *bcl2fastq --use-bases-mask Y26,I8,Y98*. Fastq quality was assessed with multiqc^85^ (version 1.9). Fastq files were aligned to the GRCh38 genome with *cellranger count* (=> version 3.0.2). Doublet scores were calculated with Scrublet (version 0.2.3) with default settings on the cellranger filtered count matrices. Cells with doublet scores below 0.5 were accepted for downstream analysis performed with scanpy^86^ (version 1.7.2, functions abbreviated with *sc* from here on) in python (version 3.7). All replicates were loaded and merged using the *sc*.*concatenate* function. *<u>Cell QC:</u>* Violin and scatter plots of QC parameter distributions were manually inspected for all samples and the following joint criteria were applied: (i) at least 100 features per cell, (ii) a maximum mitochondrial gene fraction of 20%, (iii) a maximum ribosomal gene fraction of 40%, and (iv) a maximum of 30,000 counts. Transcripts were accepted if present in at least three cells. All replicates were jointly total-count normalized excluding the top 10% highly expressed genes, logarithmized (*X*=log(*X*+1), natural logarithm), and reduced to 5000 highly variable genes. The data was scaled to a maximum value of 5.

### Cell cycle scoring

Cell cycle scores (S-, G2M-Score) and cell cycle phase (S, G2M, G1) were assigned based on a previously published cell cycle defining gene list^87^ with the *sc.score_genes_cell_cycle* function.

### Dimensionality reduction, embedding, clustering

Normalized, log-transformed and scaled data were objected to calculation of principal component analysis (PCA) coordinates, loadings, and variance via *sc.tl.pca*. The neighborhood graph was calculated using the first 50 principal components and embedded utilizing the Uniform Manifold Approximation and Projection (UMAP)^88^ algorithm via *sc.api.tl.umap*. Clusters were identified with the Leiden algorithm^89^ at resolution 0.1. Coarsenesses lower than the default parameters were chosen to reproduce approximately five cell types experimentally validated in HLOs^27^. Clusters were required to represent all individual replicates. The force-directed graph was computed via *sc.tl.daw_graph* implementing ForceAtlas2^90^ with default parameters.

### Cluster marker gene identification and pathway enrichment analysis

Cluster-characterizing genes were defined using scanpy’s implementation of the Wilcoxon rank-sum test (comparing each cluster against all other clusters) followed by Benjamini–Hochberg correction (*sc.tl.rank_genes_group*).

### Batch correction and force-directed graph drawing

Batch effects were corrected with the scanpy implementation of Harmony^91^ applying the *sce.pp.harmony_integrate* function followed by recalculation of the neighborhood graph and UMAP based on the Harmony representation. Convergence was reached prior to the maximum of 10 iterations.

### Cell cluster annotation

*<u>Automated cluster annotation based on literature markers.</u>* Literature research was performed to assemble a set of specific marker genes for cell types potentially emerging in HLOs, including mature liver cells and progenitors (Supplementary Table 1). The top 200 Wilcoxon rank sum test derived marker genes of each cluster were forwarded to the *gseapy.enrichr*^92,93^ function together with the literature reference list, the number of genes in the dataset was provided as the background parameter. Results from *enrichr* with an adjusted *p*-value below 0.05 were assigned as the new cluster identity based on the number of significantly enriched marker genes. If *p*-value criteria were not met, the label with the lowest adjusted *p*-value and maximum number of matching genes was chosen as the identity. Accepting the label “Embryonic stem cells“ required expression of either *SOX2, NANOG, POU5F1* or *KLF1*. *<u>Automated annotation based on curated databases.</u>* The ScType^40^ marker gene set (accessed 2022-Nov-25, http://session.asuscomm.com/database.php) was used as a reference for the statistical enrichment described above. Cholangiocyte-like cells were further sub-clustered and annotated as described. Hepatocyte-like cells were sub-clustered and annotated to previously published marker genes from a single-cell atlas of human liver development^51^ (pairwise hepatocyte DGE from “Source Data Fig. 1”, accessed 2022-Dec-10: https://static-content.springer.com/esm/art%3A10.1038%2Fs41556-022-00989-7/MediaObjects/41556_2022_989_MOESM4_ESM.xlsx, sheet “Hepatocyte_pairwise DEG”). Genes with a negative log fold change in HB2 compared to HB1 were used as HB1 marker genes since no HB1-specific pairwise genes with a positive log fold change were available, for all other hepatocyte stages genes with a positive log fold change compared to their respective precursor state were selected as marker genes, and cells were annotated as described above. *<u>Cell type specific pathway enrichment analysis.</u>* For each cell type the top 600-700 genes identified by the Wilcoxon rank sum test were forwarded to EnrichR^92^/gseapy^93^ (version 0.10.7, *“WikiPathways_2021_Human“* gene set, organism *“Human“*).

### Hepatocyte zonation score

Marker genes for six available adult human hepatocyte clusters were retrieved from adult human livers scRNAseq data^51^ (cluster DGE from “Source Data Fig. 1”, accessed 2022-Dec-10, https://static-content.springer.com/esm/art%3A10.1038%2Fs41556-022-00989-7/MediaObjects/41556_2022_989_MOESM4_ESM.xlsx), reduced to the top 100 genes per cluster and sub-clustered cells were annotated at as described above. Next, each hepatocyte cluster was assigned an integer between 0 and 5 (Periportal-C5:0, Periportal-C14:1, Periportal-C6:2, Interzonal-C15:3, Pericentral-C1:4, Pericentral-C3:5) reflecting the position of the cluster on the periportal-to-pericentral spectrum defined by the authors of the reference study^51^, and this value constituted the zonation score.

### SingleCellNet annotation

*<u>Generation of a reference data object.</u>* For comparing HLO cells to human liver cells, human 10X scRNA-seq data were chosen as a reference^41^. Respective *cellranger count* outputs were downloaded via: wget ftp://ftp.ncbi.nlm.nih.gov/geo/series/GSE156nnn/GSE156625/suppl/GSE156625%5FHCCFgenes%2Etsv%2Egz; wget ftp://ftp.ncbi.nlm.nih.gov/geo/series/GSE156nnn/GSE156625/suppl/GSE156625%5FHCCFmatrix%2Emtx%2Egz; wget ftp://ftp.ncbi.nlm.nih.gov/geo/series/GSE156nnn/GSE156625/suppl/GSE156625%5FHCCFbarcodes%2Etsv%2Egz and loaded into scanpy via *sc.read_10x_mtx* to obtain an *AnnData* object with raw count data. To add cell types defined by the authors, the public *HCCF1F2.h5ad* file of the respective dataset was downloaded from: https://data.mendeley.com/public-files/datasets/6wmzcskt6k/files/8ecdfe82-955a-40bd-b6b7-50edd67e9d31/file_downloaded and relevant metadata columns (*“louvain”, “NormalvsTumor”*) were extracted from the *adata.obs* dataframe, integrated into the previously generated *AnnData* object, and were added to the raw count data composing the new *AnnData* object. *<u>SingleCellNet classification of HLO-cell types based on human reference data.</u>* Using pysinglecellnet (version 0.1), the created *AnnData* object was defined as the reference dataset. A classifier was trained with the top 15 genes and gene pairs, respectively. Genes were not limited to highly variable genes. Subsequently, each HLO sample was classified. To this end, each cellranger filtered feature matrix raw output (*matrix.mtx, features.tsv, barcodes.tsv*) was loaded into scanpy as an individual query sample and underwent classification. The annotated cell type was stored as a categorical named *“SCN_class”* in the *adata.obs* slot from where it could be retrieved for downstream analyses.

### Inflammatory gene and fibrosis scoring

To render gene lists for biological processes related to fibrotic and inflammatory injury scenarios, publicly available GO term^54^ gene lists were browsed via AmiGO2^94^ filtering for Homo sapiens genes only and the *“bioentity_label”* column was exported. All GO terms are listed in Supplementary Table 4. Duplicate values and genes not present in the *adata.var* slot were removed from the lists prior to scoring Finally, *sc.tl.score_genes* with the number of control genes set to the size of the input gene list was used for scoring cells.

### DGE and pathway enrichment analysis

*<u>Differential gene expression (DGE).</u>* DGE was performed on raw count data for each cluster separately in pairwise comparisons between treatments and respective controls (e.g. Control^TGF-β1^ vs. TGF-β1) applying the Wilcoxon rank-sum test after exclusion of mitochondrial and ribosomal genes. *<u>Pathway enrichment analysis.</u>* For each cluster a hypergeometric enrichment test was performed on the top 200 differentially expressed genes using EnrichR^92^/gseapy^93^ (version 0.10.7, *“WikiPathways_2021_Human“* gene set, organism *“Human“*).

### CellPhoneDB cell-cell interactome

CellPhoneDB^56^ (version 2.1.7) with rpy2 (version 3.0.5) was run on *h5ad* files containing log-transformed count data to infer cell-cell interactions. Metadata from previously harmony-integrated and cell-type annotated *AnnData* objects served as the input to the *statistical_analysis* function along with *“gene_name“* set as an identifier for genes in the counts data. To generate the count network data and inputs for dotplots and heatmaps, *plot_heatmap_plot* and *plot_dotplot* functions were applied, and plots were created in python (version 3.7). The fraction of total interactions was calculated by dividing each cell-cell interaction count by the sum of all interactions in the respective condition group. The delta value of interactions between treatment and control condition was calculated by subtracting the fraction for each cell-cell interaction in the treatment group by the corresponding fraction of the cell-cell interaction in the control group. Clustered heat maps with euclidean distance dendrograms were created using seaborn^95^ based on either the delta of interactions or the total interaction count. Chord diagrams were computed with the python packages holoviews (version 1.15.3) and bokeh (version 3.0.3) based on the delta of interactions and computed separately for interactions with a positive delta value (labeled “enforced”) or a negative delta value (labeled “reduced”) with respect to the control condition. Mean expression indicated by dot size in all dot plots refers to CellPhoneDB output *“means“*, and is the mean of the respective two individual partners‘ average expression. Venn diagrams displaying overlapping and distinct receptor-ligand interactions, the CellPhoneDB output file *“pvalues”* served as an input. The list was filtered for significant interactions (*p*-value < 0.05), and each interaction received an identifier for its cell-cell-pair and receptor-ligand pair (e.g. “NRP1_VEGFA_Cholangiocytes|Hepatic stellate cells”). Next, the overlapping and distinct fractions were calculated based on lists of identifiers found in each treatment condition (Supplementary Table 6), and Venn diagrams generated using *pyvenn* (Adam Labadorf, https://github.com/adamlabadorf/pyvenn). For representing induced receptor-ligand pairs the set was filtered to receptor-ligand pairs with an increase of at least two significant cell-cell pairs in the treatment condition compared to the control.

### Palantir trajectory inference

ForceAtlas2-generated matrices served as distances for Palantir^65^ trajectory analysis. For lineage subsetting, the whole AnnData object was reduced to hepatocyte-, cholangiocyte- and DC-like cells for the hepatic progenitor lineage. The HSC-lineage included all HSC-, SMC- and fibroblast-like cells. Early cells for the hepatic progenitor lineage were automatically defined by searching for the cell with the highest *AFP* expression among all cells expressing *CEBPA*. For the HSC lineage, the cell with the highest *IGFBP2* expression among all cells expressing *DDIT4* was chosen, based on reference data of fetal HSCs^51^. No terminal or start cells were predefined. Palantir analysis was performed with 30 nearest-neighbors, 30 components and 9000 numerical waypoints.

### Retrieval of human NAFLD signatures in injured HLOs

Previously published 26 and 98 gene signatures predicting fibrosis in NAFLD^68^ were transferred to lists, and each cell from the integrated dataset of all orbital-shaker cultured HLOs (*n* = 8 controls, *n* = 2 OA, *n* = 2 PA, *n* = 2 TGF-β1, total 82,467 cells, quality control and processing as described above) obtained a score for the respective gene lists as described for other inflammatory and fibrosis-associated scores. Dot plots were generated with scanpy based on scaled count data for the entire dataset or for each cell type individually splitting into the treatment groups. For statistical analysis, lists of numerical score values of all cells from each group (control, OA, PA, TGF-β1), either in the full dataset or per cell cluster and were forwarded to statistical testing described below.

### Statistics and reproducibility

All statistical tests were performed in python (version 3.6) with scipy.stats^96^ (version 1.8.0) and scikit-posthocs^97^ (version 0.6.7). Two groups were compared using the two-tailed Mann-Whitney U test followed by a Bonferroni correction. For three or more groups, the two-tailed Kruskal-Wallis non-parametric test followed by a post hoc Conover’s test with Bonferroni correction was applied. A difference of mean between groups was considered significant at a *p*-value below 0.05. *N* numbers for individual HLO samples in qPCR and immunohistochemistry experiments indicate individual experiments (individual differentiation cycles starting from day 0 hPSCs). For scRNA-seq, *n* of samples refer to individual HLO single cell suspensions that arose from the same differentiation experiment if performed in replicate, e.g. for day 21 HLOs 3 replicates (*n* = 3) were sequenced, which in this case means from one HLO experiment, single cell suspensions from three different wells from the same treatment and time point were harvested and underwent individual library preps.

### Data availability

Raw data for single-cell RNA sequencing are available at Gene Expression Omnibus (GEO) under the accession GSE207889. Previously published scRNA-seq data that were re-analysed can be found at GSE156625 and GSE130073.

### Code availability

The following custom scripts are available upon request and will be made publicly available upon release.

## Supplementary information

Supplementary Information includes 7 figures and 8 tables:

### Supplementary Figures

|  |  |
| --- | --- |
| **Supplementary Figure 1** | Representative images of fixed HLOs. Corresponding to Figure 1c |
| **Supplementary Figure 2** | Replicate distribution, marker gene expression, and SingleCellNet<sup>42</sup> annotation for 10X scRNA-seq data of day 21 ULA-HLOs. Corresponding to Fig. 1d |
| **Supplementary Figure 3** | Induction of fibrotic phenotypes (morphology, qPCR, Sirius red staining) in HLOs treated with OA, PA, and TGF-β1. Corresponding to Fig. 2 |
| **Supplementary Figure 4** | Marker gene expression and replicate distribution in cells from control and injured HLOs. Corresponding to Fig. 4b |
| **Supplementary Figure 5** | Pathway enrichment analysis, cell-type resolved. Corresponding to Fig. 4f |
| **Supplementary Figure 6** | Differential cell-cell interactions in OS-HLOs upon OA, PA, and TGF-β1 treatment. Corresponding to Fig. 5 |
| **Supplementary Figure 7** | Cell type-resolved expression of genes predicting fibrosis stages in NAFLD. Corresponding to Fig. 7c |

### Supplementary Tables

|  |  |
| --- | --- |
| **Supplementary Table 1** | Marker gene sets used in this study, related to Figure 1 |
| **Supplementary Table 2** | Ranked cluster-defining genes in ULA-HLOs and OS-HLOs, related to Figures 1 and 3 |
| **Supplementary Table 3** | Cell type proportions for control, PA, OA, and TGF-β1-treated HLOs, related to Figure 4 |
| **Supplementary Table 4** | GO terms for scoring inflammation and fibrosis, related to Figure 4 |
| **Supplementary Table 5** | DGE for all cell types for PA, OA, and TGF-β1-treated HLOs, related to Figure 4 |
| **Supplementary Table 6** | Significant ($p < 0.05$) receptor-ligand interactions from CellPhoneDB analysis of cell-cell interactions for PA, OA, and TGF-β1-treated HLOs, related to Figure 5 |
| **Supplementary Table 7** | Inflammation and fibrosis-related genes ordered by imputed expression at final pseudotime for stellate and hepatocyte HLO lineages, related to Figure 6g |
| **Supplementary Table 8** | Sequences of human qPCR primers used in this study |

## Acknowledgments

The authors thank Gary Bader, Tallulah Andrews, and Ramnik Xavier for helpful discussions, Cristin McCabe and Jacques Deguine for assistance with data management, and Adam Slamin, Dan Dubinsky, and the Broad Genomics Platform for help with generation of single cell sequencing data. The authors also thank the Massachusetts General Hospital (MGH) NextGen Sequencing Core for additional assistance with generation of single cell sequencing data and Annika Gabriel for designing color palettes used in the manuscript.

